# PIP30/FAM192A is a novel regulator of the nuclear proteasome activator PA28γ

**DOI:** 10.1101/160739

**Authors:** Beata Jonik-Nowak, Thomas Menneteau, Didier Fesquet, Véronique Baldin, Catherine Bonne-Andrea, Francisca Méchali, Bertrand Fabre, Prisca Boisguerin, Sylvain de Rossi, Corinne Henriquet, Martine Pugnière, Manuelle Ducoux-Petit, Odile Burlet-Schiltz, Angus I. Lamond, Philippe Fort, Séverine Boulon, Marie-Pierre Bousquet, Olivier Coux

## Abstract

PA28γ is a nuclear activator of the 20S proteasome involved in the regulation of several essential cellular processes, such as cell proliferation, apoptosis, nuclear dynamics and cellular stress response. Unlike the 19S regulator of the proteasome, which specifically recognizes ubiquitylated proteins, PA28γ promotes the degradation of several substrates by the proteasome in an ATP- and ubiquitin-independent manner. However its exact mechanisms of action are unclear and likely to involve additional partners that remain to be identified. Here we report the identification of the first cofactor of PA28γ, PIP30/FAM192A. PIP30 binds directly and specifically via its C-terminal end and in an interaction stabilized by casein kinase 2 phosphorylation to both free and 20S proteasome-associated PA28γ. Its recruitment to proteasome-containing complexes depends on PA28γ and its expression increases the association of PA28γ with the 20S proteasome in cells. Further dissection of its possible roles shows that PIP30 alters PA28γ-dependent activation of peptide degradation by the 20S proteasome *in vitro* and negatively controls in cells the presence of PA28γ in Cajal Bodies by inhibition of its association with the key Cajal body component coilin. Altogether, our data show that PIP30 deeply affects PA28γ interactions with cellular proteins, including 20S proteasome, demonstrating that it is an important regulator of PA28γ in cells and thus a new player in the control of the multiple functions of the proteasome within the nucleus.

**Significance Statement:** The 20S proteasome is a key actor of the control of protein levels and integrity in cells. To perform its multiple functions, it works with a series of regulators, among which a nuclear complex called PA28γ. In particular, PA28γ participates in the regulation of cell proliferation and nuclear dynamics. We describe here the characterization of a novel protein, PIP30/FAM192A, which binds tightly to PA28γ and favors its interaction with the 20S proteasome while inhibiting its association with coilin, a central component of nuclear Cajal bodies. Thus PIP30/FAM192A critically controls the interactome and consequently the functions of PA28γ, and appears to be a new player in the fine regulation of intracellular proteostasis in the cell nucleus.

## Introduction

The 20S proteasome is a multimeric “barrel-like” protease that bears multiple peptidase (i.e., trypsin-like, caspase-like and chymotrypsin-like) activities inside its internal catalytic chamber (1). It plays a major role in the regulated degradation of intracellular proteins and requires for its functioning, in most cases, the binding of regulatory complexes to one or both of its ends (1). Among these regulators, the well-characterized 19S regulatory complex has been shown to associate with the 20S core to form the 26S proteasome that specifically recognizes and degrades polyubiquitylated proteins (2, 3). Another type of 20S proteasome regulators is the PA28 (also called 11S or REG) family of complexes, which comprises in higher eukaryotes three related proteins (PA28α, -β and -γ) that associate in two distinct complexes (4). PA28α and PA28β form a cytoplasmic heteroheptamer, PA28αβ (5, 6), which participates in the immune response by favoring the generation of MHC class I antigens (7, 8). In contrast, PA28γ is essentially nuclear and forms a homoheptamer (9, 10). PA28 complexes mediate primarily ubiquitin-independent proteolysis when bound to the 20S proteasome. However, they are also found in hybrid proteasomes, i.e. 20S proteasome bound to a PA28 heptamer on one end and a 19S complex on the other end (11) and are thus likely to also participate in ATP-dependent degradation of ubiquitylated proteins.

Although PA28γ has been little studied, it is involved in various essential cellular pathways, including cell proliferation, through the degradation of the cell cycle inhibitors p21^Cip1^, p16^INK4A^ and p19^Arf^ (12–14). PA28γ also participates in the Mdm2-dependent degradation of the tumor suppressor p53 (15). Furthermore, PA28γ^-/-^ mice display growth retardation (16) and PA28γ is overexpressed in several types of cancer (17–19). In addition to its involvement in the control of cell proliferation, PA28γ is implicated in the regulation of chromosomal stability (20) and plays important roles in nuclear dynamics by modulating the number and size of various nuclear bodies, including Cajal bodies (CBs) (21), nuclear speckles (22) and PML bodies (23). It is also involved in the cellular stress response, as it is recruited to double strand break (DSBs) sites upon DNA damage (24) and is required for the UVC-induced dispersion of CBs (21).

Despite its important cellular functions, the mechanisms of action of PA28γ, like those of the related PA28αβ complex, remain elusive. It is known that, as the 19S complex, PA28 complexes open the gated pore of the 20S proteasome’s α-ring upon binding, thus allowing easier substrate entrance into the catalytic chamber (25). Artificial peptide substrates commonly used to measure proteasome activities are believed to passively diffuse through the 20S pores, and therefore PA28s’ stimulation of their degradation can be explained by 20S-pore opening only. However, how PA28 complexes recruit protein substrates and deliver them to the proteasome is not understood, as these complexes are *a priori* inert molecules that, unlike the 19S complex, do not possess any ATPase activity that could provide energy and movement to unfold the substrates and inject them into the 20S proteasome (2, 3). A likely possibility is thus that PA28 complexes function in proteasome-dependent proteolysis in association with other proteins that remain to be defined.

In this study, we describe the first partner of a PA28 complex, the evolutionary conserved PIP30/FAM192A protein. We show that PIP30 binds with high specificity to PA28γ and enhances its association with the 20S proteasome in cells. Importantly also, PIP30 binding affects PA28γ specificity towards peptide substrates *in vitro* as well as its interactions with cellular proteins such as the CB marker coilin. Therefore, PIP30 is a major regulator of PA28γ functions and consequently of the nuclear functions of the proteasome.

## Experimental procedures

### Phylogenetic analyses

Genomes were explored by using Annotation and BLAST search tools available in the Geneious 9.1.7 software package (http://www.geneious.com/). Amino acid sequences were aligned (MAFFT v7.017) and phylogenetic tree was deduced by maximum likelihood analysis (PhyML).

### Antibodies

Antibodies and related agents used in this study are described in supplemental Material and Methods.

### Production of 20S proteasome and recombinant proteins

Native 20S proteasome was purified from extracts of HeLa cells (IPRACELL, Belgium) using classical chromatographic procedures (26). Unless indicated, recombinant PIP30 was produced in *E. coli* as a 6His-tagged protein and purified by affinity purification followed by proteolytic removal of the tag, anion exchange chromatography and gel filtration (Fig. S8B). At the last step, the protein was eluted slightly earlier than the BSA (67kDa) marker, i.e. at an apparent molecular weight larger than twice what was expected. Human recombinant PA28αβ was produced and purified from *E. coli*, as previously described (27). Human PA28γ was expressed in *E. coli* BL21 DE3 CodonPlus. After expression, the PA28γ complex was purified by chromatography as PA28αβ, with some minor modifications of the procedure. Both PA28αβ and PA28γ complexes efficiently activated the peptidase activities of the 20S proteasome. Recombinant GST and GST-PIP30-H201 fusion proteins were produced in bacteria and efficiently purified using glutathione sepharose beads.

### Pulldown and immunoprecipitation

For immunoprecipitation of GFP-fusion proteins, U2OS cells were transfected with the indicated constructs. Twenty-four hours post-transfection, cells were homogenized in lysis buffer (25 mM HEPES pH 7.8, 100 mM KCl, 10 mM MgCl_2_, 1 mM EDTA, 1% IGEPAL CA-630, 0.1% Triton X-100, 1 mM DTT, 1 mM ATP, 10% glycerol (v/v)), in the presence of complete EDTA free protease inhibitor cocktail (Roche), for 15 min on ice. After centrifugation at 15000xg for 15 min (4°C), supernatants were recovered and protein concentration was determined by Bradford assays using BSA as a standard. 20 μl of GFP-TRAP-A® beads (Chromotek) were used per IP, mixed with 200 μg of protein extract and incubated with constant gentle stirring for 1 h at 4°C.

Beads were washed three times with lysis buffer and boiled in 2X Laemmli sample buffer. Samples were then analyzed by SDS-PAGE and immunoblotting.

Endogenous PIP30 and PA28γ were immunoprecipitated from total cell extracts using anti-PIP30 (rabbit) and anti-PA28γ (rabbit) antibodies, bound to protein A magnetic beads (Dynal, Lake Success, NY) for 2 h at 4°C.

For co-immunoprecipitation of coilin and PA28γ proteins (Fig 5B), nuclear extracts were prepared as described in (21). Briefly, U2OS and PIP30^-/-^ cell pellets were resuspended in hypotonic buffer (10 mM Tris-HCl pH 7.5, 10% glycerol, 4 mM DTT, 50 mM NaF, 1 mM Na_3_VO_4_, 1 mM MgCl_2_) in the presence of complete EDTA-free protease inhibitor cocktail (Roche). IGEPAL CA-630 was then added at the final concentration of 0.5% and cells were incubated on ice for 3 min. After centrifugation at 800xg for 5 min, the nuclei, present in the pellet, were resuspended in digestion buffer (2 mM Tris-HCl pH 8.5, 20% glycerol, 10 mM DTT, 50 mM NaF, 1 mM Na_3_VO_4_, 1 mM MgCl_2_, 5 mM CaCl_2_, 1X complete protease inhibitor cocktail) supplemented with 75 U/ml micrococcal nuclease, and then digested for 15 min at 25°C with constant stirring. At the end, an equivalent volume of extraction buffer (2 mM Tris-HCl pH 8.5, 50 mM NaF, 1 mM Na_3_VO_4_, 1 mM MgCl_2_, 20 mM EDTA, 0.84 M KCl, 1X complete protease inhibitor cocktail) was added and the mix was incubated on ice for 20 min. Nuclear extracts were clarified by centrifugation for 30 min at 15000xg. Before immunoprecipitation, KCl concentration was reduced to 280 mM, 3 μg of anti-coilin or control IgG were added to 400 μg of nuclear extracts and incubated for 2 h at 4°C. Immunoprecipitated proteins were collected by addition of 15 μl of protein A-sepharose beads. After extensive washes, beads were boiled in 2X Laemmli sample buffer and samples were analyzed by SDS-PAGE and immunoblotting.

### Mass spectrometry analyses

SILAC IPs (endogenous PA28γ and GFP-FAM192A/PIP30) were essentially performed as previously described in (28). Further details are provided in supplemental section.

Whole human proteasome complexes, including 20S-bound activators and regulators, were immunopurified and analyzed by quantitative mass spectrometry as previously described (29, 30). 2×10^8^ *in vivo* formaldehyde-crosslinked human cells (HeLa and U937, 3 biological replicates per cell line) were used. For complete nuclear proteasome interactome analysis, U937 cells nuclei were prepared and, before proteasome purification, the purity of nuclear proteins was assessed both by western blot and MS analysis, as detailed earlier (31). Purified proteasome complexes were analyzed by mass spectrometry as previously described (32). Further details are provided in supplemental section.

### Native electrophoresis in Tris-glycine system

Recombinant protein samples were incubated 5-10 min at room temperature (RT) in reaction buffer (Tris-HCl 20mM, pH 7.5, DTT 1mM, Glycerol 10% (v/v)), then supplemented with 1µL of native sample buffer (xylene cyanol FF in reaction buffer supplemented with 50% glycerol) and applied on 5% polyacrylamide gel prepared in Tris-Glycine electrophoresis buffer (25mM Tris-HCl pH 8.0, 192mM Glycine, 1mM DTT). Native electrophoresis was performed for 4-5hrs (100V, 4^°^C). After denaturation in 10X TG-SDS buffer, proteins were transferred on PVDF membrane and immunoblotted.

### Surface Plasmon Resonance analysis

Experiments were performed on Biacore 3000 apparatus (GEHealthcare) at 25°C using a flow rate of 50 µl/min in HBS-EP buffer (GE Healthcare). To compare the binding of PA28γ and PA28αβ on 6His-PIP30 recombinant protein, 6His-PIP30 (4500RU)) was captured on anti–HisTag covalently immobilized on a CM5 sensor chip using amine coupling procedure according to the manufacturer’s instructions. A control flowcell was obtained with the same chemical procedure without protein. 60µl of PA28γ and PA28αβ (50µg/ml) were injected on His6PIP30 and control flowcells followed by a dissociation step of 400s.

### *In vitro* CK2 phosphorylation assay

GST and GST-PIP30-H201 proteins were incubated with recombinant CK2 according to the manufacturer’s instructions. For radioactive kinase assays, ^33^P labeled ATP (1 μCi in the presence of 100 μM cold ATP) was included. Reactions were stopped either by adding 5 mM EDTA or Laemmli sample buffer.

### Proteasome peptidase assays

Peptidase activities of 20S proteasome were measured using black flat-bottom 96-well plates (Nunc), in a final volume of 50µL, in reaction buffer (20mM Tris-HCl pH 7.5, 50mM NaCl, 1mM DTT, 10% glycerol (v/v)) supplemented with 100μM peptide substrates. When indicated, purified recombinant PA28 and/or PIP30 were added. Kinetic analyses showed that the assays are linear at least for 30 min. Peptide degradation was measured by the fluorescence emitted by the AMC group released by cleavage of the substrate (excitation 380nm, emission 440nm) using a FLx800 microplate fluorescence reader (Bio-Tek Instruments).

### Immunofluorescence microscopy and PLA assays

Cells were fixed in 3.7% paraformaldehyde/PBS for 10 min at RT, washed with PBS and permeabilized in PBS containing 1% Triton X-100 for 15 min at RT. Coverslips were then blocked in blocking solution (1% FCS, 0.01% Tween-20/PBS) for 10-20 min and incubated with primary antibodies, diluted in blocking buffer, for 1hr at RT or 37°C in a humidified atmosphere. After three washes in PBS, coverslips were incubated with Alexa-Fluor conjugated secondary antibodies, diluted in blocking solution for 40 min at RT. Coverslips were washed with PBS, incubated with 0.1 μg/ml DAPI solution in PBS for 5 min at RT, washed twice in PBS and finally once in H_2_O. Coverslips were mounted on glass slides using ProLong Gold antifade reagent (Thermo Fisher Scientific).

For proximity ligation assays (PLA), cells on coverslips were fixed and permeabilized as described in (22). Coverslips were then blocked in a solution provided by the Duolink® kit. Cells were then incubated with mentioned antibodies as described above. Duolink® *In Situ* PLA Probe Anti-Rabbit MINUS and Anti-Mouse PLUS and Duolink® *In Situ* Detection Reagents (Sigma-Aldrich) were used, according to the manufacturer’s instructions.

### Image acquisition and analysis

Z-stacks and images were acquired with a 63X/1.4 NA or 40X oil immersion objective lenses using widefield microscopes, DM6000 (Leica Microsystems) or Axioimager Z1 or Z2 (Carl Zeiss), equipped with coolSNAP HQ2 cameras (Photometrics). Images were acquired as TIF files using MetaMorph imaging software (Molecular Devices). For CB and PLA dots quantitative analysis, Z-stacks were acquired every 0.3 to 0.4μm (Z step) with a range of 10-15 μm to image the whole nuclei. The number of PLA foci was analyzed with ImageJ. The size, the intensity and the number of Cajal bodies were measured with ImageJ (1.49v), using a specific “macro” that has been created to automatically quantify these different parameters. The script allows creating a mask of DAPI image to isolate the nucleus of each cell and create a maximum intensity projection (MIP) of the 25 Z-stacks. The mask is used in the MIP to count the number of CB or PLA dots of each nucleus via an appropriate thresholding. The “Analyze Particles” tool of ImageJ was used to calculate the size and the mean gray value of each Cajal body.

## Results

### FAM192A is a novel partner of both free and 20S proteasome-bound PA28γ that regulates the interaction between PA28γ and 20S proteasome

In order to identify new interactors of PA28γ, we used a high throughput approach combining endogenous PA28γ immunoprecipitation (IP) and SILAC-based quantitative proteomics (28) (Fig. S1A, left panel). In this experiment, 68 human protein groups containing at least two unique peptides were quantified and visualized by plotting log_2_(H/L ratio) versus log_2_(H intensity) (Fig. 1A, Table S1). PA28γ itself was found with a H/L SILAC ratio close to 15. Among PA28γ interactors was FAM192A, a poorly annotated nuclear protein of unknown cellular function, whose expression is induced during skeletal muscle atrophy (33). FAM192A was also identified with a high SILAC ratio in a GFP-PA28γ pull-down that we performed in parallel (Table S2). Conversely, PA28γ was the most abundant and almost unique partner pulled-down with GFP-FAM192A (Fig. 1B, Fig. S1A right panel and Table S3)). Reciprocal IPs on endogenous proteins showed that at least 70% of each protein coprecipitates with the other (Fig. S1B). Like PA28γ, FAM192A is nuclear (Fig. S8A) and interaction between the two proteins occurs within the nucleus (Fig. S1C).

**Fig. 1:**
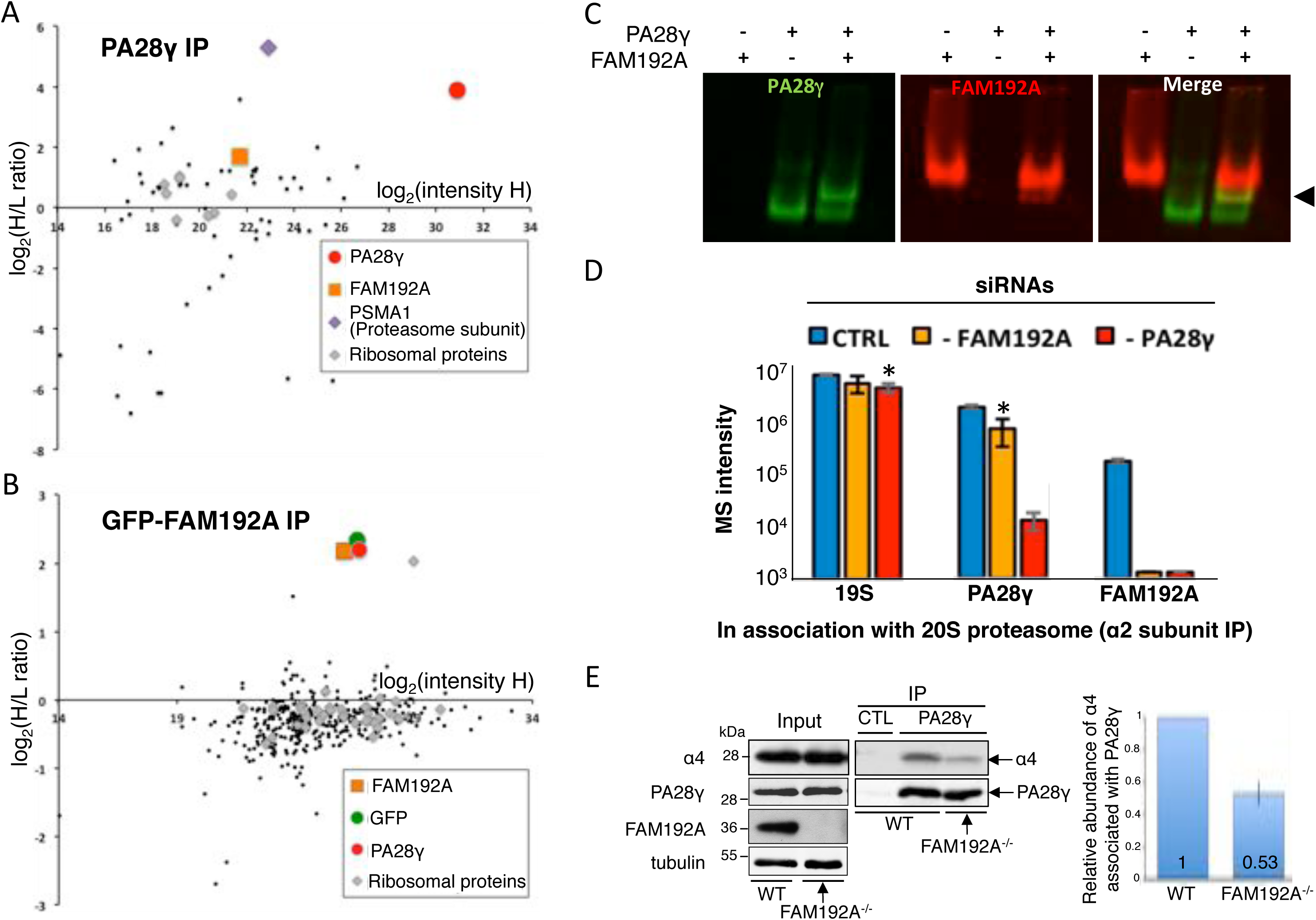
FAM192A is a novel and direct interactor of PA28γ that increases 20S proteasome/PA28γ interaction. **(A)** Graphic representation of the result of endogenous PA28γ SILAC IP in U2OS cells, visualized by plotting log_2_(H/L) versus log_2_(H intensity) values for all 68 human protein groups quantified by MaxQuant with a minimum of 2 unique peptides identified. Proteins specifically interacting with the bait, i.e., PA28γ, are expected to show Heavy/Light (H/L) ratios higher than 1. In contrast, proteins non-specifically binding to the beads (experimental contaminants) are expected to show H/L ratios close to 1. Proteins with ratios lower than 1 are mostly environmental contaminants, such as keratins. **(B)** Graphic representation of the result of the SILAC GFP-TRAP IP of GFP-FAM192A, visualized by plotting log_2_(H/L) versus log_2_(H intensity) values for all 364 protein groups quantified by MaxQuant with a minimum of 2 unique peptides identified. In both graphs, PA28γ is indicated by a red dot, FAM192A is indicated by an orange square and GFP by a green dot. The PSMA1 subunit of the 20S core proteasome is indicated by a violet diamond. Ribosomal proteins, which can be considered here as non-specific interaction partners, are indicated by grey diamonds. **(C)** Recombinant PA28γ and FAM192A directly interact with each other, as analyzed by native electrophoresis. After electrophoresis, the proteins were transferred on membrane and immunoprobed with specific antibodies visualized with an Odyssey infrared imaging system (LI-COR Biosciences). The arrowhead on the right indicates the formation of a complex (yellow) between purified PA28γ (green) and PIP30 (red). **(D)** Effect of FAM192A and/or PA28γ knock-down on the incorporation of 19S, PA28γ, and FAM192A into proteasome complexes. Proteasome complexes were immuno-purified using the MCP21 antibody (targeting the α2 subunit of 20S proteasome) from total extracts of siRNA-treated HeLa cells. Proteins were identified and quantified by nano-LC-MS/MS analysis. SD were calculated from triplicates. *: p = 0.02 **(E)** Quantification of PA28γ-bound 20S proteasome in wild-type and FAM192A^-/-^ U2OS cells, as determined by co-immunoprecipitation of α4 subunit from 150 µg of total cell extract with anti-PA28γ antibodies. The right panel is constructed from three independent experiments (error bars: SD).

Using recombinant FAM192A and PA28γ proteins produced in *E. coli*, we found that the interaction between PIP30 and PA28γ is direct, as shown for example by native gel electrophoresis (Fig. 1C) or Surface Plasmon Resonance (SPR) (Fig. S1D). SPR showed no interaction between PIP30 and the closely related PA28αβ complex (Fig. S1D), demonstrating the high specificity of PA28γ/PIP30 interaction *in vitro*.

Through parallel approaches, we investigated putative nuclear Proteasome Interacting Proteins by quantitative analysis of proteasome complexes immuno-purified from nuclei of U937 cells using the MCP21 antibody (targeting the α2 subunit of 20S proteasome), as described earlier (31). As expected, all 20S proteasome and 19S regulatory complex subunits, as well as known proteasome interactors such as PA28γ, PA200 and USP14 were found in this category (abundance ratios (IP 20S / IP control) above 2 and p-value below 0.05 (n=3)) (Fig. S2A and Table S4). Interestingly, the protein FAM192A showed an enrichment factor of 6.4 and a p-value < 5.10^-4^ and thus clearly qualifies as a new proteasome interacting protein.

Further quantitative analysis of proteasome-associated proteins after PA28γ knockdown in HeLa cells showed that FAM192A is recruited on proteasomes in a PA28γ-dependent manner (Fig.1D). The enrichment in anti-FAM192A immuno-precipitates of all subunits of the 19S regulatory complex (PSMCx and PSMDx) (Fig. S2B) demonstrated that FAM192A is also recruited, via its interaction with PA28γ, on hybrid proteasomes.

Interestingly, co-immunoprecipitation of PA28γ with 20S proteasome was significantly (p-value = 0.02) reduced in FAM192A-depleted cells (Fig. 1D, please note that the scale is in log). Co-IP of wild type and FAM192A^-/-^ U2OS cell extracts showed that FAM192A depletion elicits a 2-fold decrease in PA28γ/α4 interaction (Fig.1E). Thus FAM192A promotes the association of PA28γ with the 20S proteasome.

Altogether, our experiments establish that FAM192A is a major direct interactor of proteasome-free and proteasome-bound PA28γ and that it favors the association of PA28γ with the 20S proteasome. FAM192A is also called NIP30 (NEFA-Interacting Protein 30kD) and/or might refer to a NIP (4-hydroxy-5-iodo-3-nitrophenyl acetate) – labeled 30kD protein that interacts with phosphatidylinositol biphosphate (34). Based on the above experiments, and since it has no clear assigned cellular function, we propose to rename it PIP30, for PA28γ Interacting Protein 30 kDa. This name will be used thereafter in this manuscript.

### PA28γ / PIP30 interaction is stabilized by CK2 phosphorylation of PIP30 C-terminus

Evolutionary analyses show that PIP30 sequence is characterized by the presence of three conserved domains: an N-terminal signature domain (pfam10187, also called NIP30 domain, ≈100 amino acids) and two smaller motifs (Fig. S3A). The pfam10187 domain is found in metazoa as well as in all major eukaryotic supergoups. This allows tracing back PIP30 gene to early stages of eukaryote evolution (Fig. S3B), as it is the case for PA28γ (4). The conserved C-terminal motifs of PIP30 proteins are enriched in phosphorylatable residues, including a completely conserved tyrosine and several serine residues.

Using various truncation mutants, we determined that the last 54 amino acids of PIP30 (aa 201-254, i.e., H201 mutant), are necessary and sufficient for interaction with endogenous PA28γ and that aa 223-230 are critical for the association (Fig. 2A and S4A). This C-terminal region includes the highly conserved serine-rich and acidic sequence highlighted in Fig. S3A and contains two SDSE motifs (Fig. S4B) that both match the canonical consensus site (S/T-X-X-E/D/pS) for casein kinase 2 (CK2) (35, 36). Indeed, we found that this region can be phosphorylated by CK2 *in vitro* (Fig. 2B). Phosphorylation is dependent on the integrity of the CK2 consensus sites, as it was reduced when the serine residues S_222_ and S_228_ were replaced by alanine (mutant SS-AA), and abolished or strongly impaired when the acidic residues within the two CK2 consensus motifs were converted into basic lysine residues (mutants D_223_E_225_-KK and D_229_E_231_-KK, respectively) (Fig. 2B and S4B).

**Fig.2:**
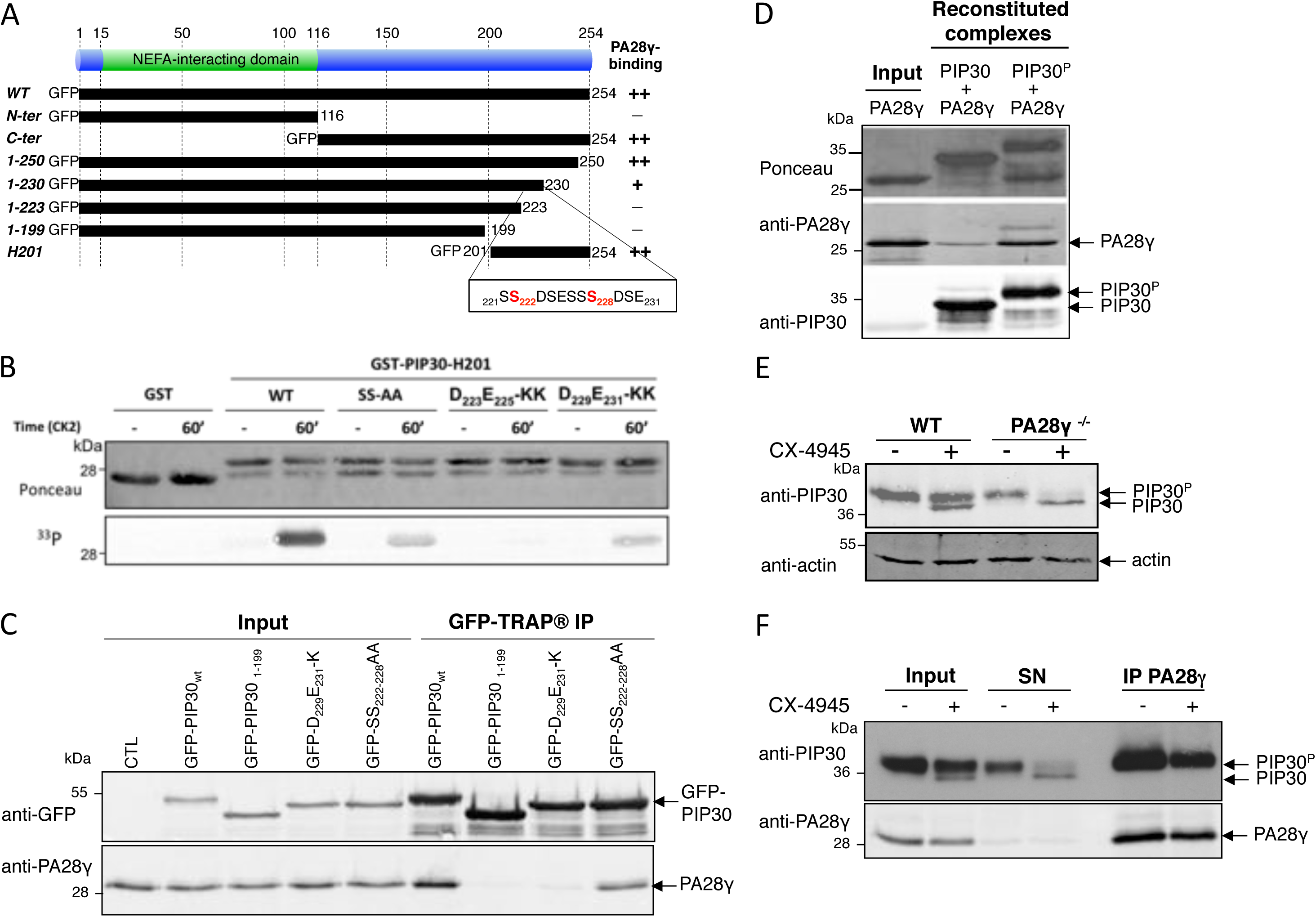
PIP30 interacts with PA28γ via its CK2-phosphorylated C-terminus. **(A)** Graphical summary of the analysis of the interaction between several GFP-PIP30 truncation mutants and PA28γ. **(B)** The C-terminal end of PIP30 (GST-PIP30-H201) is phosphorylated *in vitro* by CK2 and point mutations in its CK2 consensus sites strongly reduce its phosphorylation. Purified GST, GST-PIP30-H201 and GST-PIP30-H201 mutants were incubated with CK2, in the presence of [^33^P]-ATP for the indicated times. The presence of GST-tagged proteins was revealed by Ponceau staining and their phosphorylation was detected by autoradiography. **(C)** Point mutations in the CK2 consensus motifs of GFP-PIP30 strongly reduce the interaction between GFP-PIP30 and PA28γ in cells. U2OS cells were transiently transfected with wild type or the indicated mutants of PIP30 fused to GFP. GFP-tagged proteins were pulled-down using GFP-TRAP® beads and the presence of co-immunoprecipitated PA28γ was assessed by western blot. Wild type GFP-PIP30 was used as a positive control, GFP-PIP30_1-199_ mutant as a negative control. **(D)** PA28γ has more affinity for CK2-phosphorylated PIP30 (PIP30^p^) than for unphosphorylated PIP30 *in vitro*. Bacterially expressed and purified His-PIP30 was either phosphorylated, or not, by CK2 *in vitro* and then mixed with recombinant PA28γ. The complex was purified on a Ni-NTA column and analyzed by SDS-PAGE and western blot with anti-His anti-PA28γ antibodies. **(E)** The association between PA28γ and PIP30 protects PIP30 from dephosphorylation. Wild type and PA28γ^-/-^ U2OS cells were either treated, or not, with CX-4945 (10 µM) for 24 h and the PIP30 PAGE migration profile was analyzed by western blot. The phosphorylated form (PIP30^P^) and the unphosphorylated form of PIP30 are indicated. **(F)** PA28γ interacts with the phosphorylated form of endogenous PIP30 in cells. Endogenous PA28γ was immunoprecipitated from 500 μg of total extracts from U2OS cells either treated, or not, with CX-4945 (10 µM) for 24 h. The presence of co-immunoprecipitated PIP30 was assessed by western blot. The input and unbound (SN) fractions were also analyzed. After CX-4945 treatment, two bands appear for endogenous PIP30, the phosphorylated form (PIP30^P^) and the unphosphorylated form (PIP30), as indicated.

In the context of the full length GFP-PIP30 protein, these mutations altered the binding to PA28γ in cells. In a manner parallel to their effects on CK2 phosphorylation *in vitro*, the acidic mutants abolished or strongly impaired the binding while the SS-AA mutant reduced it (Fig. 2C). The weaker binding of the SS-AA mutant strongly suggests that phosphorylation is stabilizing PA28γ/PIP30 interaction. Indeed, using purified proteins, PA28γ showed more affinity for CK2-phosphorylated than for non-phosphorylated PIP30 (Fig. 2D).

To verify that CK2 is the endogenous kinase phosphorylating PIP30, we treated cells with the selective CK2 inhibitor CX-4945 (37). This treatment revealed a second, faster-migrating band (Fig. 2E), suggesting that in cells PIP30 is primarily in a CK2-phosphorylated form. While only a minor fraction of endogenous PIP30 was non-phosphorylated in wild type cells upon CK2 inhibition for 24hrs, it was mainly non- or hypo-phosphorylated in PA28γ^-/-^ cells after the same treatment (Fig. 2E). This suggests that, although PIP30 does not require PA28γ for being phosphorylated, the binding to PA28γ protects it from being dephosphorylated. This was confirmed by the fact that λ-phosphatase can dephosphorylate a fraction of immunoprecipitated endogenous PIP30, but not the endogenous PIP30 co-immunoprecipitated with PA28γ (Fig. S4C). Finally, after CX-4945 treatment, only the phosphorylated PIP30 was retrieved upon PA28γ immunoprecipitation (Fig. 2F), showing that the phosphorylation of PIP30 by CK2 in cells stabilizes its interaction with PA28γ. Altogether our results show that the C-terminal end of PIP30 protein is critical for its binding to PA28γ and that its phosphorylation by CK2 stabilizes this interaction.

### PIP30 controls substrate diffusion to the catalytic chamber of the proteasome

We next assessed whether PIP30 could interfere with the best-known property of PA28γ, i.e. its ability to activate the peptidase activities of the proteasome *in vitro*.

Using recombinant PIP30 in combination with PA28γ and 20S proteasome, we found that PIP30 differentially altered degradation of a panel of standard proteasome peptide substrates (38) by the PA28γ-activated proteasome (Fig. 3A). For example, while the activation of Suc-LLVY-amc (LLVY) and Ac-nLPnLD-amc (nLPnLD) degradation was partially inhibited, degradation of Boc-LRR-amc (LRR) was almost insensitive to the presence of PIP30 (Fig. 3A and S5A). PIP30 had no effect when PA28αβ was used to activate the 20S proteasome in similar experiments (Fig. S5B). This indicates that PIP30 is not directly affecting 20S proteasome and most likely acts in this assay through its interaction with PA28γ. The differential effects of PIP30 were not correlated with specific proteasome peptidase activities (Fig.3A), suggesting that PIP30 is not acting through alteration of the proteasome active sites. To confirm this hypothesis, we used the proteasome activity probe Bodipy TMR-Ahx_3_L_3_VS (a generous gift of Prof. Huib Ovaa), which labels efficiently the β2 (trypsin-like activity) and β5 (chymotrypsin-like activity) subunits of the proteasome (39, 40). We found that the ratio between β2 and β5 labeling was not altered by PIP30 (Fig.3B), showing that the relative activity of both sites is not changed. Therefore, the differential degradation of LRR (β2 substrate) and LLVY (β5 substrate) elicited by PIP30 is most likely due to a differential diffusion rate into the catalytic chamber of the proteasome. This strongly suggests that PIP30 regulates peptide entrance or transit through the channel opened by PA28γ upon binding to the 20S proteasome.

**Fig.3:**
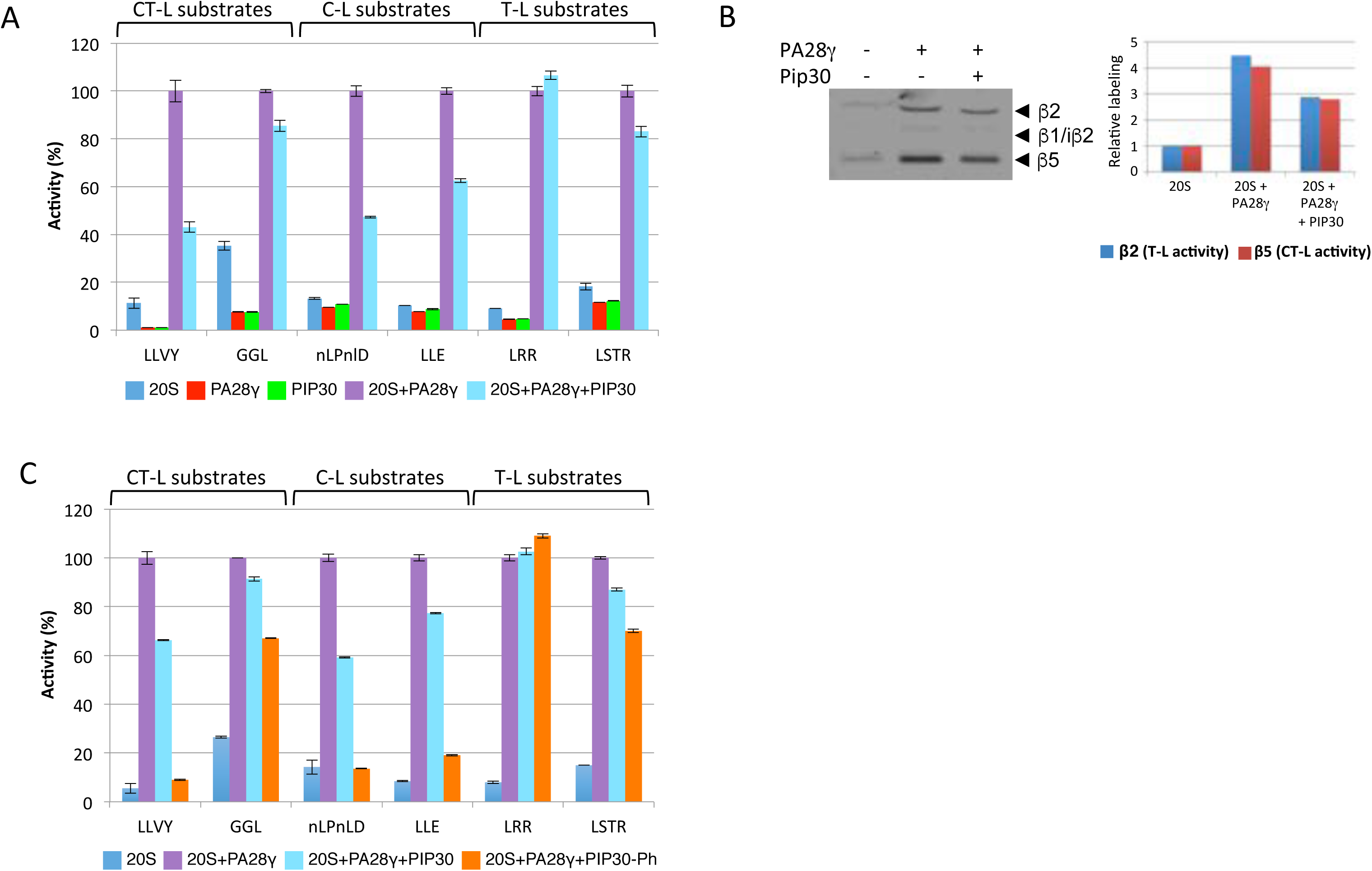
PIP30 differentially affects peptide diffusion towards the catalytic chamber of the proteasome. **(A)** Inhibition by PIP30 of PA28γ-activated 20S proteasome is peptide-substrate specific. Proteasome activity was assayed in the presence of the indicated peptides and proteins and normalized by setting to 100% the value obtained in the presence of 20S proteasome and PA28γ. The assays were performed for 20 min at 37°C in a 50μl reaction mixture containing the indicated combinations of 20S proteasome (0.5µg), PA28γ (1µg), and PIP30 (4µg). The respective final concentrations of the 20S proteasome and PA28γ are set up such as the peptidase activities are not fully activated (i.e. PA28γ is limiting in the assay). At the final concentration used for PIP30, the inhibitory effect is maximal (Fig. S5A). Error bars represent deviation from the mean of technical duplicates. The figure is representative of >3 distinct experiments. Peptides used are substrates of the following peptidase activities of the 20S proteasome: Suc-LLVY-amc, Z-GGL-amc : chymotrypsin-like activity; Ac-nLPnLD-amc, Z-LLE-amc : caspase-like activity; Boc-LRR-amc, Boc-LSTR-amc : trypsin-like activity. **(B)** The activity of the chymotrypsin-like and trypsin-like catalytic sites of the 20S proteasome is not modified by PIP30: 20S proteasome (2 μg) was mixed in reaction buffer (Tris-HCl 20mM pH 7.5, DTT 1mM, glycerol 10%), in a final volume of 18µL, with either no protein or with PA28γ (6μg) preincubated for 5 min at RT with or without PIP30 (8.3µg). After incubation 5 min at RT, 2µL of the probe Bodipy TMR-Ahx_3_L_3_VS (1µM in reaction buffer) were added to each sample and the mixtures were further incubated 15min at 37°C. After denaturation with Laemmli buffer and electrophoresis, the labeled proteasome subunits were visualized (left panel) using a Typhoon FLA 7000 (GE Healthcare). The labeling of subunits β2 and β5 was quantified and plotted after normalization by the value of their labeling in 20S proteasome alone (right panel). In the experimental conditions used, the labeling of both subunits is roughly proportional to their activity. The figure is representative of 3 distinct experiments. **(C)** Comparison of the effects of PIP30 and its CK2-phosphorylated form (PIP30-Ph) on the peptidase activities of the PA28γ / 20S proteasome complex. The experiments were performed as in (A) except that only 2 μg of PIP30 and PIP30-Ph were used per assay. Error bars represent deviation from the mean of technical duplicates. The figure is representative of 3 distinct experiments.

We next assessed whether PIP30 phosphorylation affects its effect on peptide degradation. Comparing non-phosphorylated and CK2-phosphorylated PIP30 (Fig. S8D), we found that phosphorylation of PIP30 enhanced its inhibitory effects (Fig. 3C): For the most affected peptides, phosphorylated PIP30 was able to abrogate the activating effect of PA28γ on their hydrolysis. However, it had a limited effect on other peptides and remained without effect on the basic LRR substrate (Fig. 3C).

Overall, these data show that PIP30 is able to alter the 20S proteasome-activating properties of PA28γ *in vitro*. In this process, it seems to act as a phosphorylation-modulated filter that differentially impairs the entrance or the transit of peptide substrates through the PA28γ channel.

### PIP30 alters PA28γ functions in Cajal body dynamics

We have previously shown that PA28γ overexpression leads to the disruption of CBs (21), which are evolutionary conserved coilin-dependent and coilin-rich subnuclear compartments. CBs are involved in the maturation of small nuclear RNPs (snRNPs) and small nucleolar RNPs (snoRNPs) as well as in the processing of histone pre-mRNAs (41, 42). We thus tested whether PIP30 has also a role in CB dynamics. First, we compared CB number in wild type, PA28γ^-/-^ and PIP30^-/-^ U2OS cells. We found that while 80% of wild type and PA28γ^-/-^ U2OS cells display CBs, only 40% of PIP30^-/-^ cells are CB-positive (Fig. 4A and 4B). Furthermore, in cells displaying CBs, the absence of PIP30 leads to a decrease in the average number of CBs per nucleus (Fig.4C). The effect of PIP30 depletion was rescued by expression of GFP-PIP30, but not of GFP-PIP30_1-199_, a mutant that cannot bind to PA28γ (Fig. 4D). Together, these results show that PIP30, like PA28γ (21), controls the steady state number of CBs and that this depends on its binding to PA28γ. However, PIP30 and PA28γ do not have the same effects in this process.

**Fig. 4:**
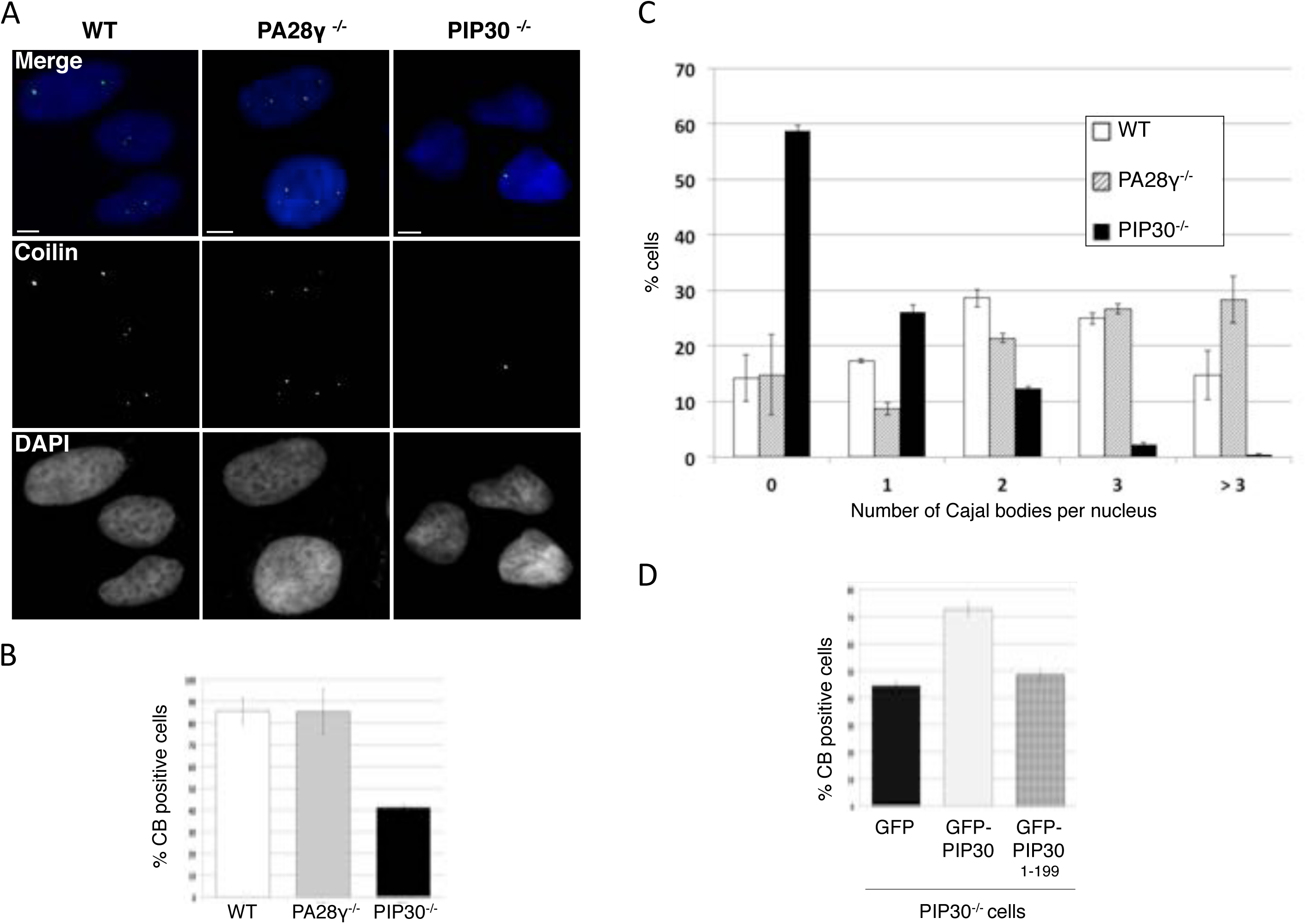
PIP30 is involved in CB dynamics. **(A)** PA28γ and PIP30 depletion have differential effects on Cajal body dynamics. The presence of CBs in wild type, PA28γ^-/-^ and PIP30^-/-^ U2OS cells was analyzed by indirect immunofluorescence using 5P10 anti-coilin antibody. Bars, 10 μm. **(B)** The coilin signal was used to determine the percentage of CB-positive cells for wild type, PA28γ^-/-^ and PIP30^-/-^ U2OS cells. Data represent the mean of two independent experiments (n= 526 WT, 532 PA28γ ^-/-^ and 503 PIP30 ^-/-^ cells). **(C)** PIP30 depletion induces a decrease in the mean number of CBs per nucleus. The coilin signal was used to quantify the number of CBs per nucleus in wild type, PA28γ^-/-^ and PIP30^-/-^ U2OS cells. Data show the percentage of cells displaying 0, 1, 2, 3 or more CBs per nucleus and represent the mean of two independent experiments (n= 526 WT, 532 PA28γ ^-/-^ and 503 PIP30 ^-/-^ cells). **(D)** The overexpression of GFP-PIP30, but not of the GFP-PIP30_1-199_ mutant defective for PA28γ binding, rescues the formation and/or stability of CBs in U2OS PIP30^-/-^ cells. PIP30^-/-^ cells were transfected with GFP, GFP-PIP30 WT and GFP-PIP30_1-199_. The percentage of CB positive cells was determined in each condition by analyzing the presence/absence of coilin dots in all GFP-positive cells. n= 63, 88, and 93 for transfected cells with GFP, GFP-PIP30, GFP-PIP30_1-199_, respectively.

In the absence of PIP30, i.e. in siRNA PIP30-depleted or PIP30 knockout cells, but not in control cells, we observed an accumulation of PA28γ in all residual CBs, either identified by coilin (Fig. 5A) or WRAP53 (another CB marker) staining (Fig. S6A). Furthermore, re-expression of GFP-PIP30 in PIP30-depleted cells abrogated the accumulation of PA28γ in CBs (Fig. S6B). These observations demonstrate that PIP30 inhibits PA28γ subnuclear localization in CBs.

Since PA28γ is known to bind to the key CB component coilin ((21) and Table S2), we analyzed whether PIP30 could affect this process. We observed that PIP30 depletion led to an increased interaction of PA28γ with coilin, as seen by co-immunoprecipitation experiments (Fig. 5B). In line with this observation, in the absence of PIP30, an increased number of dots could be seen by PLA using anti-PA28γ and coilin antibodies, supporting the notion that PIP30 inhibits PA28γ/coilin interaction (Fig. S7A and B).

**Fig. 5:**
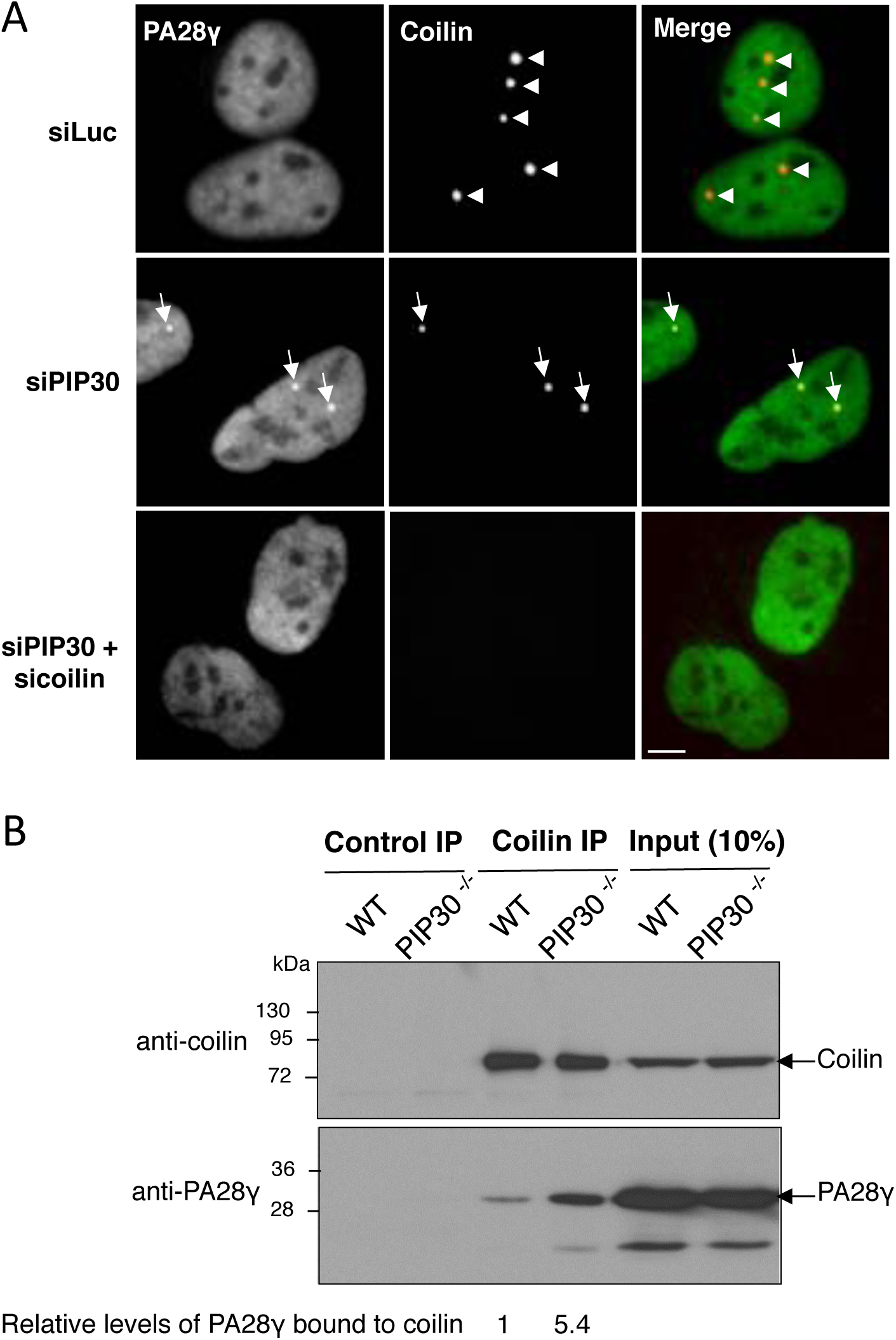
PIP30 depletion induces accumulation of PA28γ in residual Cajal bodies and enhances the interaction of PA28γ with coilin. **(A)** PIP30 depletion induces PA28γ accumulation in Cajal bodies. The co-localization of endogenous PA28γ and coilin was analyzed by indirect immunofluorescence in U2OS cells treated either with a control siRNA (siLuc), a siRNA targeting PIP30 or siRNAs targeting both PIP30 and coilin, and in PIP30^-/-^ cells. In the merge panels, the coilin signal is in red and the PA28γ signal in green. Bars, 10μm. **(B)** PIP30 depletion increases the level of endogenous coilin co-immunoprecipitated with PA28γ. Nuclear cell extracts from asynchronously growing parental (WT) or PIP30^-/-^ U2OS cells were incubated with control IgG or anti-coilin antibodies. Immunocomplexes were analyzed by SDS-PAGE, probed for the indicated proteins, and the amount of PA28γ associated with coilin was quantified and normalized to the amount of immunoprecipitated coilin. The figure is representative of 3 distinct experiments.

Altogether, these experiments show that PIP30 effect on CBs is mediated through its ability to bind PA28γ and counteract PA28γ association with coilin, confirming that PIP30 indeed functions as an endogenous regulator of PA28γ.

## Discussion

Despite the fact that PA28 complexes have long been known, little is understood regarding their roles as proteasome regulators, besides their ability to open upon binding the gated channel of the 20S proteasome. However, since specific protein substrates have been described for PA28γ, it is possible that these regulators are not just passively opening the gate of the 20S proteasome but somehow contribute to substrate selection and possibly unfolding and injection into the 20S core. If true, then it is likely that they work in synergy with other proteins, since by themselves they seem a priori unable to actively promote proteasomal degradation of specific protein substrates. We thus performed various proteomics experiments to characterize proteins associated with PA28γ.

Among the interaction partners of endogenous PA28γ, we found several proteins involved in the regulation of RNA processing (Table 1), suggesting an important role of PA28γ in this function. Although PA28γ interacts with the 20S proteasome, we identified only the α6 subunit (PSMA1) in the conditions used. However, we found all subunits with high H/L ratios when PA28γ pulldowns were performed in the presence of proteasome inhibitors (data not shown), confirming that PA28γ/20S proteasome interaction is labile (31) and is stabilized upon proteasome inhibition (11, 43).

Importantly, we identified a novel and prominent partner of PA28γ, PIP30/FAM192A. Interaction between PIP30 and PA28γ has already been listed in large-scale proteomics experiment data in drosophila and human (44, 45). However, our results are the first to validate its biological relevance. We demonstrated that this interaction is direct and occurs in cells with 20S-proteasome free and bound PA28γ. The interaction is specific for PA28γ since we found no interaction between PIP30 and PA28αβ *in vitro*. This suggests that interaction may involve the homolog-specific insert, which is the most divergent domain between PA28 paralogs (46). Importantly we show that the PA28γ-binding region of PIP30 is located in its C-terminal end and that CK2 phosphorylation of a short motif in this region stabilizes its association with PA28γ. Since mutation of serine residues (S_222_ and S_228_) of both typical CK2 targets does not completely abolish binding, it is likely that additional serine residues are phosphorylated by CK2. This is line with the ability of CK2 to catalyze the generation of phosphoserine stretches (36), and with the report that S_219_, S_221_, S_222_ and S_224_ of PIP30 are phosphorylated *in vivo* (Global Proteome Machine, Accession: ENSP00000335808 (47)). The role of these multiple phosphorylation events is not clear. CK2 is indeed an ubiquitous and pleiotropic kinase (48), and even if its activity has been shown to be positively regulated by the Wnt/β-catenin pathway (49) it is considered to be constitutively active in cells. Our results also suggest an extremely slow turnover of the phosphate groups for the majority of PIP30 bound to PA28γ, probably because steric hindrance makes them inaccessible to phosphatases once the complex is formed. It thus seems unlikely that CK2 phosphorylation of PIP30 is used in cells to dynamically control PIP30/PA28γ interaction.

Comparison of PIP30 and PA28γ distributions within the eukaryotic phylogenetic tree (Fig. S3B) shows that their presence or absence is not correlated. For example, in all Ascomycota including *S. cerevisiae,* PIP30 is present while PA28γ has been lost. This indicates that the two proteins may function independently from each other. However, in human cells, more than 70% of PA28γ and PIP30/FAM192A are associated in the same complex (Fig. S1B), suggesting that they cooperate in most of their functions.

In line with this assertion, our data show that PIP30 regulates the function of PA28γ as a proteasome regulator, since its invalidation halves the intracellular level of the PA28γ / 20S proteasome complex (Fig. 1D, E). However, since the 20S proteasome / PA28γ complex only represents ≤ 5% of the total 20S proteasome (29), it is difficult to predict the biological impact of this effect. Based on our *in vitro* analyses showing that PIP30 inhibits the degradation of some peptides by reconstituted 20S proteasome / PA28γ complexes but is permissive for others (Fig. 3), it is tempting to speculate that PIP30 acts as a molecular sieve that hinders entrance of some protein substrates into the PA28γ channel while being neutral for others. The likely binding of PIP30 to the PA28γ specific insert that forms a loop located close to the pore of the complex (25, 50) is compatible with such a molecular sieve role. Ongoing structure/functions analyses aiming at precisely mapping where PIP30 interacts with PA28γ will help to answer these questions.

A second illustration of the role of PIP30 as a PA28γ regulator is the demonstration that both proteins intimately cooperate in the regulation of CB integrity. Indeed, PIP30 depletion leads to a strong decrease in the number of CBs and to an increased interaction between coilin and PA28γ (Fig.5B, S7A,B). This mimics the phenotype induced by PA28γ overexpression ((21), Table S3). PIP30 depletion also elicits the accumulation of PA28γ in residual CBs, while PA28γ is usually not detectable in these structures. To our knowledge, PA28γ has only been observed in CBs of motor neurons from patients with Type I spinal muscular atrophy (SMA) (51). In these pathologic neurons, the assembly of CBs is impaired, due to the lack of the Survival Motor Neuron (SMN) protein, an essential CB component (52–54). Since CBs are dynamic structures undergoing constant remodeling (55), the accumulation of PA28γ in residual CBs observed in both PIP30^-/-^ cells and SMA motor neurons could reflect the fact that these residual CBs are stalled at transient intermediate stages of assembly/disassembly in which PA28γ is involved. Alternatively, the absence of PIP30 may result in the formation of defective CB structures that are not normally present in wild type cells.

Together, the effects of PIP30 deletion on CB integrity suggest that increasing the levels of PA28γ / coilin complexes negatively regulates the number of CBs and that PIP30 inhibits the association between PA28γ and coilin. This model supports the idea that PA28γ overexpression leads to CB destabilization by overwhelming PIP30 inhibition and therefore that the equilibrium between free PA28γ and PIP30-bound PA28γ is an important parameter in this process. In this regard, it is interesting to note that CB fragmentation upon UV-C treatment is associated with the concomitant increase in PA28γ and its recruitment to coilin (21). If PIP30 levels are limiting in cells, the resulting excess of PIP30-free PA28γ might be sufficient to interact with coilin and induce CB fragmentation.

Altogether, the results presented here show that PIP30 is an important partner of PA28γ that regulates its interactome, e.g. by stabilizing its interaction with the 20S proteasome and inhibiting its interaction with coilin. Since most PA28γ is bound to PIP30 in standard cell culture conditions, it is clear that PIP30 must now be taken into account when studying PA28γ. Although much remains to be understood regarding PIP30 biological functions, our results represent a significant breakthrough since they provide the first clues on the regulation of PA28γ/20S proteasome complex as well as novel angles of attack to dissect PA28γ functions and mechanisms of action.

## Acknowledgements

The authors wish to thanks Prof. Huib Ovaa (Leiden, the Netherland) for proteasome activity probes, Dr. Anne Blangy (CRBM, France) for critical reading, the MRI facility for image and data analysis. We also acknowledge the rotation students who participated to this work.

This work was supported by the CNRS, and by funding from the People Programme (Marie Curie Actions) of the EU Seventh Framework Programme FP7 to O.C. (REA agreement n°290257, « UPStream »), the Association pour la Recherche sur le Cancer to S.B. (ARC n°PJA 20141201831 and n°SFI20111203984), the French Ministry of Research (Investissements d’Avenir Program, Proteomics French Infrastructure, ANR-10-INBS-08) and the Fonds Européens de Développement Régional (FEDER), Toulouse Métropole, the Région Occitanie (fellowships to TM and BF) to O.B.-S. B.J.N was supported by the Marie Curie International Training Network « UPStream ». S.B. was initially supported by EMBO and HFSP Long Term Fellowships.

The authors declare no competing financial interests.

## Abbreviations

CB: Cajal body
CK2: Casein Kinase 2
H/L ratio: Heavy/Light ratio
IP: Immunoprecipitation
PIP30: PA28g Interacting Protein 30kDa
PLA: Proximity Ligation Assay
SILAC: Stable Isotope Labeling by Amino-Acids in Cell Culture
U2OS: human OsteoSarcoma cell line
LLVY: Suc-LLVY-amc
GGL: Z-GGL-amc
nLPnLD: Ac-nLPnLD-amc
LLE: Z-LLE-amc
LRR: Boc-LRR-amc
STLR: Boc-LSTR-amc-amc, amido-4-methylcoumarin
MG132: Z-Leu-Leu-Leu-al

## Author contributions

OBS, AIL, MPB, SB and OC designed research; BJN, TM, DF, VB, CBA, FM, BF, PB, CH, MP, MDP, PF, MPB, SB and OC performed research; SdR contributed new analytical tool; BJN, TM, DF, VB, CBA, FM, BF, PB, CH, MP, MDP, OBS, AIL, PF, MPB, SB and OC analyzed data; PF, MPB, SB and OC wrote the paper

## Supplemental information

### Material and Methods

#### Cell culture and transfections

U2OS, HEK293T, HCT116, RKO, and HeLa cell lines (ATCC) were grown in DMEM medium (Lonza) containing 4.5mg/ml glucose and supplemented with 10% inactivated FBS (ThermoScientific). U937, HeLa S3, and NB4 cell lines were grown in RPMI 1640 media supplemented with 10% FBS. KG1a cell line was grown in RPMI 1640 media supplemented with 20% FBS. MRC5 cell line was grown in MEM-a media supplemented with 10% FBS. All cell lines were cultured with 2mM L-glutamine (Lonza), 100U/ml penicillin and 100µg/ml streptomycine (Lonza) in humid atmosphere containing 5% CO_2_ at 37°C.

Cells were typically transfected with 20nM siRNA and 0.5 µg/ml DNA, using Lipofectamine RNAiMAX (Thermo Fisher Scientific) or Dharmafect (Dharmacon) and Jet-PEI (Ozyme) transfection reagents respectively, according to manufacturer’s instructions. To generate stable GFP-PIP30/FAM192A and GFP-PA28γ U2OS cell lines, parental U2OS cells were transfected with peGFP-C1 PIP30/FAM192A or peGFP-C1 PA28γ plasmids and positive clones were selected in G418-containing medium.

#### Antibodies and other reagents

Unless otherwise specified, the sources of the antibodies used in this study were as follows: anti-PA28γ (rabbit polyclonal, BML-PW8190, ENZO Life Sciences; rabbit polyclonal, 383900, Zymed/Life Technologies; rabbit polyclonal, PD003, MBL; mouse monoclonal, 611180, BD Transduction), anti-α2 subunit of the proteasome 20S (mouse monoclonal MCP21), anti-α4 subunit of the proteasome 20S (mouse monoclonal PW8120, ENZO Life Science) anti-GFP (rabbit polyclonal, TP401, Torrey Pines Biolabs), GFP-TRAP® beads (Chromotek), anti-coilin (mouse monoclonal 5P10 (56)) for indirect immunofluorescence, rabbit polyclonal PLA 0290, Sigma-Aldrich, for PLA and IPs, rabbit polyclonal R288 (57) for WB), anti-β-actin (rabbit monoclonal, 13E5, Cell Signalling), anti-α-tubulin (mouse monoclonal, T9026, Sigma-Aldrich), anti-WRAP53 (A301-442A, Bethyl). The anti-PIP30/FAM192A rabbit polyclonal antibody was raised against full-length protein (UniProtKB - Q9GZU8) and affinity-purified. Fluorescent secondary antibodies conjugated either to AlexaFluor 488, 594 and 633, or to DyLight 680 and 800, were purchased from Molecular Probes and ThermoFisher Scientific, respectively. Secondary antibodies conjugated to HRP were purchased from BioRad. Recombinant Casein Kinase 2 was purchased from New England Biolabs (P6010S) and CK2 inhibitor CX-4945 from Selleckchem (S2248).

#### Plasmids and gene disruption

The sequence coding for human PIP30/FAM192A (Q9GZU8) was fused to that of GFP by inserting it using the HindIII and BamH1 sites in pEGFP-C1 plasmid (Clontech). GFP-PIP30 truncation mutants and point mutation mutants were generated by site-directed mutagenesis (Site-directed mutagenesis kit, Invitrogen) and verified by sequencing. For expression in *E. coli*, the cDNA coding for human PA28γ (P61289) was subcloned between the NcoI and XhoI sites of the pET14b plasmid. Fam192A/PIP30 cDNA was subcloned into the pHisTEV30a plasmid (gift from R. Hay’s group), resulting into a fusion protein with a 6His tag that can be removed by TEV digestion.

For Cas9-mediated gene disruption, guide RNAs targeting PIP30/FAM192A (GCCTCTTACCATGTTTTCTGAGG) and PSME3/PA28γ (GGAAGTGAAGCTCAAGGTAGCGG) were selected using ChopChop (https://chopchop.rc.fas.harvard.edu/index.php) and corresponding oligonucleotides were subcloned in pMLM3636 (gift from Keith Joung, Addgene, plasmid # 43860) and pUC57-U6 (gift from Edouard Bertrand’s laboratory). For PIP30 depletion, 800 bp PIP30 right and left DNA homology arms were obtained by gene synthesis (IDT, USA) and subcloned in HR110PA-1 vector (SBI, Palo Alto, USA). U2OS PIP30^-/-^ cells were generated by co-transfection of PIP30 sgGuide, pJDS246-WT CAS9 (gift from Keith Joung, Addgene, plasmid # 43861) and PIP30 HR110PA-1 vector using Turbofect (Thermofischer) (58). PSME3/PA28γ^-/-^ cells were generated by cotransfection of PSME3/PA28γ sgGuide and pX459 vectors. Both PIP30 and PSME3/PA28γ^-/-^ cells were selected with puromycine (1 µg/ml). Single clones were then expanded and analyzed by western blotting using PIP30 and PA28γ antibodies. Of note, the KO cell lines, made using the CAS9/CRISPR gene editing technology (Fig. S8C) did not show significant proliferation defects, as assessed by video-microscopy and FACS analysis (data not shown). Thus PIP30, as established previously for PA28γ, is also not essential for cell viability in normal growth conditions and does not accumulate at specific cell-cycle stage.

#### RNA interference

Silencing RNAs were obtained from Dharmacon: On target plus Human PSME3 siRNA 5nmol (J-012133-05-0005) and On target plus SMARTpool Human NIP30 5 nmol (L-014528-01-0005). Silencing RNA targeting coilin was obtained from IDT (AGAGTCGAGAGAACAATA).

#### Proteomics analyses

For SILAC IPs (28) (endogenous PA28γ and GFP-PIP30/FAM192A), parental U2OS and stable GFP-PIP30 U2OS cells were grown in DMEM medium for SILAC (ThermoFisher, 89985) supplemented with 10% dialyzed fetal calf serum (FCS) and penicillin/streptomycin for 10 days. L-arginine (R0) (84 mg/ml, Sigma-Aldrich) and L-lysine (K0) (146 mg/ml, Sigma-Aldrich) were added to the light (L) medium, and L-^13^C_6_-arginine (R6) and L-4,4,5,5-D_4_-lysine (K4) (Cambridge Isotope Laboratories) were added to the heavy (H) medium. Ten 140-mm diameter culture dishes were used per SILAC condition. Nuclear extracts were prepared by incubating cells in buffer A (20 mM Tris-HCl pH 7.4, 10 mM KCl, 3 mM MgCl_2_, 0.1% IGEPAL CA-630, 10% glycerol, protease inhibitor cocktail) for 10 min at 4°C, with occasional mixing. Nuclei were then centrifugated at 1430xg for 10 min at 4°C, resuspended in S1 solution (0.25 M sucrose, 10 mM MgCl_2_, protease inhibitor cocktail) and layered over a cushion of S3 solution (0.88 M sucrose, 0.5 mM MgCl_2_, protease inhibitor cocktail). After centrifugation at 2800xg for 10 min at 4°C, a cleaner nuclear pellet was obtained, which was resuspended in RIPA buffer, sonicated 5 x 10 s on ice, to ensure release of as many nuclear proteins as possible, and centrifuged at 2800xg for 10 min at 4°C to pellet debris.

Prior to endogenous PA28γ IP, control rabbit IgG and anti-PA28γ antibodies (rabbit polyclonal ZYMED and MBL) were covalently coupled to protein G-dynabeads (Thermo Fisher Scientific). Nuclear extracts were precleared by incubation with protein G-dynabeads alone and separate IPs were then performed in parallel by incubating the same amount of proteins from L and H extracts on control and anti-PA28γ-beads, respectively. For GFP-PA28γ and GFP-PIP30 IPs, extracts were precleared with protein G-sepharose beads alone and separate IPs were then performed by incubating the same amount of proteins from L and H extracts with GFP-TRAP® beads affinity matrix (Chromotek) for 1h at 4°C. After several washes, bound proteins were eluted in 1% SDS, boiled for 10 min, reduced and alkylated and then separated by SDS-PAGE. After tryptic *in gel* digestion, peptides were analyzed by LC-MS/MS (LTQ-Orbitrap XL, Thermo Fisher Scientific Inc.). Data were then analyzed and quantified by MaxQuant (version 1.0.12.31) (59) and the Mascot search engine (Matrix Science, version 2.2.2) software, as described in (28).

For purified proteasome complexes mass spectrometry analyses, protein eluates were precipitated with trichloroacetic acid, heated in Laemmli buffer (95°C, 30 min), and loaded on a 12% SDS-PAGE gel. For one-shot analysis of the entire mixture, no fractionation was performed, and the electrophoretic migration was stopped as soon as the protein sample entered the separating gel. The gel was briefly stained with Coomassie blue and a single band, containing the whole sample, was cut and processed. For shotgun analysis, electrophoretic migration was performed in order to fractionate the protein eluate into 10 gel bands. Proteins were in-gel digested with trypsin and the resulting peptides mixtures were analyzed by nano-LC-MS/MS using an UltiMate 3000 system (Dionex) coupled to a LTQ-Orbitrap Velos mass spectrometer (Thermo Fisher Scientific, Bremen, Germany). Five microliters of each peptide sample corresponding to an equivalent initial quantity of proteins of 2.5 µg were loaded on a C18 precolumn (300 μm inner diameter × 5 mm; Dionex) at 20 µl/min in 5% acetonitrile, 0.05% trifluoroacetic acid. After 5 min of desalting, the precolumn was switched online with the analytical C18 column (75 μm inner diameter × 15 cm; PepMap C18, Dionex) equilibrated in 95% solvent A (5% acetonitrile, 0.2% formic acid) and 5% solvent B (80% acetonitrile, 0.2% formic acid). Peptides were eluted using a 5–50% gradient of solvent B during 160 min at a 300 nl/min flow rate. The mass spectrometer was operated in data-dependent acquisition mode. Detailed MS operation settings and data analyses are described elsewhere (30).

## Legends of supplemental figures

**Fig. S1. (A)** Design of the SILAC IP experiments. For both endogenous PA28γ IP and GFP-FAM192A IP, control IPs were performed with nuclear extracts prepared from U2OS cells grown in “light”, i.e., unlabeled (R_0_K_0_) medium. Endogenous PA28γ IP was performed with a nuclear extract prepared from U2OS cells grown in “heavy” (R_6_K_4_) medium. GFP-TRAP IP of GFP-FAM192A was performed with a nuclear extract prepared from U2OS cells stably expressing GFP-FAM192A and grown in “heavy” (R_6_K_4_) medium. After the immunoprecipitation steps, eluted proteins were in-gel digested with trypsin and peptides analyzed by Liquid Chromatography Tandem Mass Spectrometry (LC-MS/MS) and quantified using MaxQuant. **(B)** U2OS cell extracts were subjected to immunoprecipitation (IP) with anti-PA28γ, anti-FAM192A and control antibodies. The relative abundance of FAM192A and PA28γ in the extract before (Input) and after IP (SN) was quantified with ImageJ software and normalized to Tubulin. **(C)** FAM192A interacts with PA28γ in the nucleus. Proximity ligation assay (PLA) was performed on U2OS cells with mouse anti-PA28γ and rabbit affinity purified anti-FAM192A primary antibodies. Green dots reveal places where PA28γ and FAM192A are in close proximity and most likely in the same complex (top panel). As a negative control, coverslips with only anti-FAM192A antibody were used (bottom panel). **(D)** Surface Plasmon Resonance (SPR) binding analysis of PA28γ and PA28αβ to His_6_-PIP30 captured by anti-His antibodies on a CM5 sensor chip. The inset displays the curves after subtraction of the His_6_-PIP30 dissociation from anti-His. Only PA28γ shows a binding response of 200 RU (resonance unit).

**Fig. S2. (A)** Volcano plot representation of 20S proteasome interactors showing that FAM192A is a new nuclear 20S Proteasome Interacting Protein. Proteasome complexes were immuno-purified from U937 cells nuclei, as described in (31). After fractionation of the protein eluate in 10 gel bands, quantitative comparison of proteins identified in the purification using the MCP21 antibody (targeting the α2 subunit of 20S proteasome) and in a control purification was used to highlight specific Proteasome Interacting Proteins. After shotgun analysis of nuclear proteasome complexes using label-free quantitative mass spectrometry, proteins exhibiting enrichment abundance ratios (IP assay/IP control) over 2 and p-value under 0.05 (n=3) can be considered as putative Proteasome Interacting Proteins. **(B)** Ranking of Proteasome subunits found in anti-FAM192A IPs, based on label-free MS quantification (log scale). FAM192A was immunopurified from whole HeLa cell lysates using anti-NIP30 (V16) (Sc-137631, Santa-Cruz biotechnology) antibodies and the co-immunopurified proteins were sorted according to their iBAQ values obtained by LC-MS/MS analysis and data bioinformatic processing using the MaxQuant software.

**Fig. S3. (A)** Conservation of PIP30 sequence across evolution **(B)** Phylogenetic analysis showing that PIP30, like PA28γ, was present in early eukaryotes. The green checkboxes and the red crosses indicate supergroups in which PIP30 and PA28 genes are present or absent, respectively.

**Fig. S4. (A)** Analysis of the interaction between GFP-PIP30 truncation mutants and PA28γ. GFP-TRAP IP experiments were performed in U2OS cells to immunoprecipitate the overexpressed GFP-PIP30 truncation mutants illustrated in Fig. 2A, and the presence of co-precipitated endogenous PA28γ was analyzed by western-blot. **(B)** The 221-SSDSESSSDSE-231 serine-rich region of PIP30 contains two typical consensus sites for CK2. Sequences of SS-AA, D_223_E_225_-KK and D_229_E_231_-KK PIP30 mutants are indicated, with the mutated residues appearing in red (SS-AA) and in green (D_223_E_225_-KK and D_229_E_231_-KK). **(C)** PA28γ-bound PIP30 cannot be dephosphorylated by λ-phosphatase. Whole cell extracts from WT U2OS cells were incubated with magnetic beads coupled with either control IgG, anti-PIP30 or anti-PA28γ antibodies. After several washes, the beads were either incubated, or not, with λ-phosphatase, as indicated. The complexes bound to the beads were then eluted and analyzed by western blot, as well as the input (10%) and the supernatants (SN) (10%), using anti-PIP30 (red) and anti-PA28γ (green) antibodies. Two bands appear for PIP30, which correspond to the phosphorylated form (PIP30^P^) and the non-phosphorylated form (PIP30) of PIP30, as indicated.

**Fig S5. (A)** Dose-dependency of the effects of PIP30 on the degradation by the PA28γ-20S proteasome complex of the peptides Suc-LLVY-amc, Ac-nLPnLD-amc, Boc-LRR-amc. 0.5μg of purified 20S proteasome and 1μg of purified PA28γ were incubated in the microplate fluorescence reader at 37°C for 30 min in a final volume of 50μl, in the presence of the indicated amounts of purified PIP30 and of 100μM of the indicated peptides. Fluorescence was read every 2 min and the slopes were used to calculate activity of each sample. The results are expressed in fold activation (compared to 20S proteasome alone), normalized by setting to 100% the value obtained for each peptide in the absence of PIP30. **(B)** PIP30 partially inhibits 20S activation by PA28γ but not by PA28αβ. The effect of PIP30 on the activation of the 20S proteasome by PA28γ or PA28αβ was measured essentially as in figure 3, using the peptide Suc-LLVY-AMC (100µM) and 0.8μg of purified 20S proteasome, 1μg of purified PA28γ or PA28αβ and 1.4 μg of purified PIP30 in a final volume of 100μl. The data were normalized by setting to 100% the value obtained in the presence of 20S proteasome and the indicated PA28. Error bars represent deviation from the mean of technical duplicates.

**Fig. S6. (A)** PIP30 depletion induces the co-localization of PA28γ with WRAP53, a component of Cajal bodies. U2OS cells were treated with either a control siRNA (siLuc) or a siRNA targeting PIP30, for 48 h. The co-localization of PA28γ and WRAP53 in these cells was then analyzed by immunofluorescence using anti-PA28γ (green) and anti-WRAP53 (red) antibodies. **(B)** Overexpression of wild-type GFP-PIP30 abolishes the recruitment of PA28γ to CBs in PIP30^-/-^ U2OS cells. PIP30^-/-^ cells were either transiently transfected, or not, with a plasmid expressing GFP-PIP30. The co-localization of PA28γ and coilin was analyzed by immunofluorescence using anti-PA28γ (red) and anti-coilin (yellow) antibodies. The presence of GFP-PIP30 in transfected cells is visualized in green. The DAPI signal is visualized in blue. The profile plots (right panels), display the pixel intensities of the yellow (green plot), the red and the blue signals along the lines present within the merge images. Bars, 10 μm.

**Fig. S7. (A)** PIP30 depletion leads to an increased interaction between PA28γ and coilin by PLA. To examine endogenous interactions between PA28γ and coilin, asynchronously growing parental (WT) or PIP30^-/-^ U2OS cells were subjected to an *in situ* proximity ligation assay (PLA) using anti-PA28γ (mouse) and anti-coilin (rabbit) antibodies. Bars, 10 µm. The figure is representative of 3 distinct experiments. **(B**) The number of PLA dots was quantified using the ImageJ software. Data represent the mean of two independent experiments (n= 123 WT cells and 112 PIP30^-/-^ cells). (**C**) PLA experiment showing no interaction between PIP30 and coilin. Endogenous interactions between PIP30 and coilin were examined in asynchronous growing U2OS cells by a PLA approach, using anti-PIP30 (rabbit) and anti-coilin (mouse) antibodies. Bars, 10 µm.

**Fig. S8. Characterization of tools: (A)** Validation of the polyclonal anti-PIP30/FAM192A antibody. Cell extracts from U2OS cells treated with either control or PIP30/FAM192A siRNAs for 48 h were analyzed by western blotting (left panel) and indirect immunofluorescence (right panel), using indicated antibodies. **(B**) Purification scheme of recombinant PIP30. The right panels show the last step of purification (Superdex 75) with the OD_280nm_ profile (top) and the SDS-PAGE analysis of the fractions of interest (bottom). **(C)** *Left panel:* Validation of PA28γ^-/-^ cells. The expression level of PA28γ was analyzed in WT and PA28γ^-/-^ U2OS cells by western blot using anti-PA28γ *Right panel:* Validation of PIP30^-/-^ cells. The expression level of PIP30 was analyzed in WT and PIP30^-/-^ U2OS cells by western blot using anti-PIP30 antibodies. Cl1 and Cl2 are two different clones selected for PIP30^-/-^ cells. In both panels anti-tubulin was used as loading control. **(D)** SDS-PAGE analysis of PIP30 and PIP30 phosphorylated *in vitro* by CK2 (PIP30^P^).

**Table. S1: List of the PA28**γ **interaction partners identified in endogenous PA28**γ **SILAC IP** Only proteins identified with a significant H/L SILAC ratio (H/L ratio > 1.4) and containing at least two unique peptides identified are listed in this table. Proteins are classified by descending order for the H/L ratio.

**Table. S2: List of the PA28**γ **interaction partners identified in GFP-PA28**γ **SILAC IP.** Only proteins identified with a significant H/L SILAC ratio (H/L ratio > 1.4) and containing at least two unique peptides identified are listed in this table. Proteins are classified by descending order for the H/L ratio.

**Table S3: List of the PA28γ interaction partners identified in GFP-FAM192A SILAC IP.** Only proteins identified with a significant H/L SILAC ratio (H/L ratio > 1.4) and containing at least two unique peptides identified are listed in this table. Proteins are classified by descending order for the H/L ratio.

**Table S4 : List of Proteasome-Interacting-Proteins identified in anti-proteasome pull-down experiments from U937 nuclear extracts (**Fig. 1C**) :** GN : Gene name; AC: Accession number; ID: Identifiant (in the UniProt Database); Description: Description of the Protein Entry; Best Id Score in Band # : Gel band where the protein displays the highest score of identification; Max Score: Maximum score of identification; Coverage (%) : % of the protein sequence covered; #QPep: Number of specific peptides used for the quantification; T-Test: Student T-test (3 biological replicates); Enrichment Factor: ratio of protein abundance in the proteasome purification compared to the control purification (mean of 3 biological replicates)

