## Supplementary Materials for "PIP30/FAM192A is a novel regulator of the nuclear proteasome activator PA28γ"

*Figure S1*

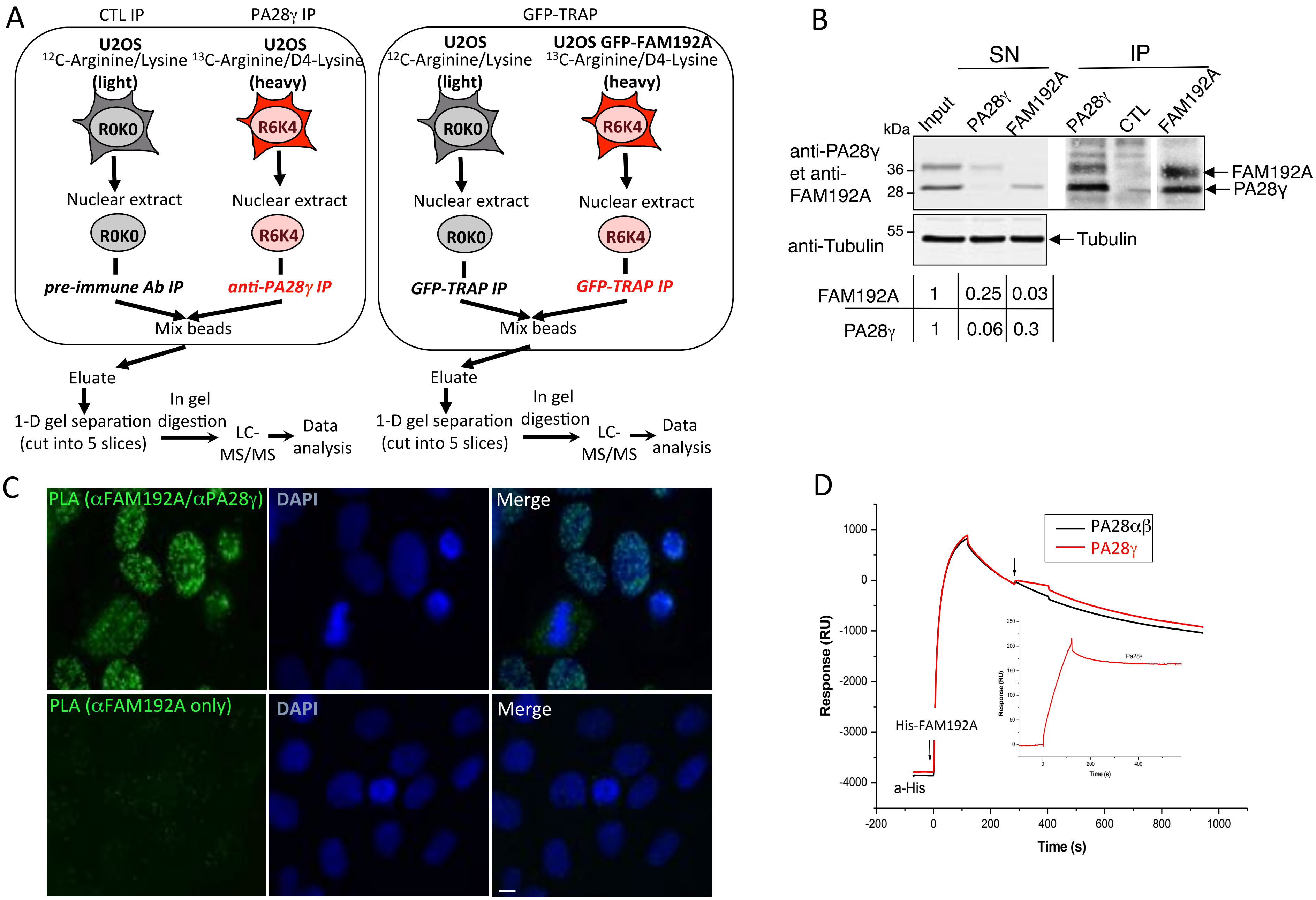
A
CTL IP
PA28γ IP
GFP-TRAP
U2OS
12C-Arginine/Lysine
(light)
U2OS
13C-Arginine/D4-Lysine
(heavy)
U2OS GFP-FAM192A
R0K0
R6K4
Nuclear extract
pre-immune Ab IP
anti-PA28γ IP
GFP-TRAP IP
Mix beads
Eluate
1-D gel separation
(cut into 5 slices)
In gel
digestion
LC-
MS/MS
Data
analysis
B
SN
IP
kDa
Input
PA28γ
FAM192A
CTL
anti-PA28γ
et anti-
FAM192A
36
28
55
anti-Tubulin
FAM192A
PA28γ
Tubulin
FAM192A 1 0.25 0.03
PA28γ 1 0.06 0.3
C
PLA (αFAM192A/αPA28γ)
DAPI
Merge
PLA (αFAM192A only)
D
PA28αβ
PA28γ
His-FAM192A
a-His
Response (RU)
Time (s)
1000
0
-1000
-2000
-3000
-4000
-200
200
400
600
800
1000

*Figure S2*

A

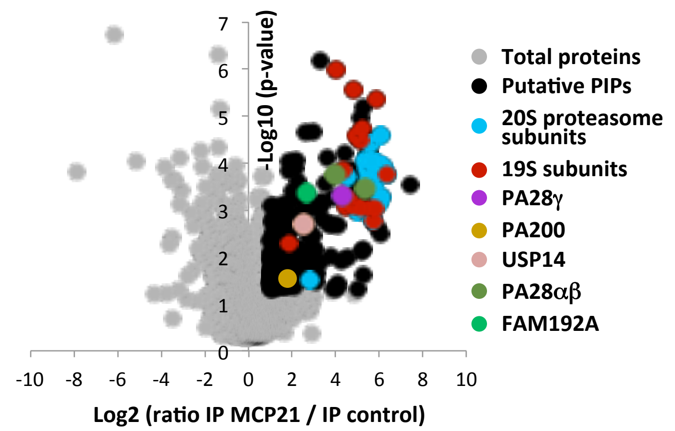

-Log10 (p-value)
Log2 (ratio IP MCP21 / IP control)
Total proteins
Putative PIPs
20S proteasome subunits
19S subunits
PA28γ
PA200
USP14
PA28αβ
FAM192A


B

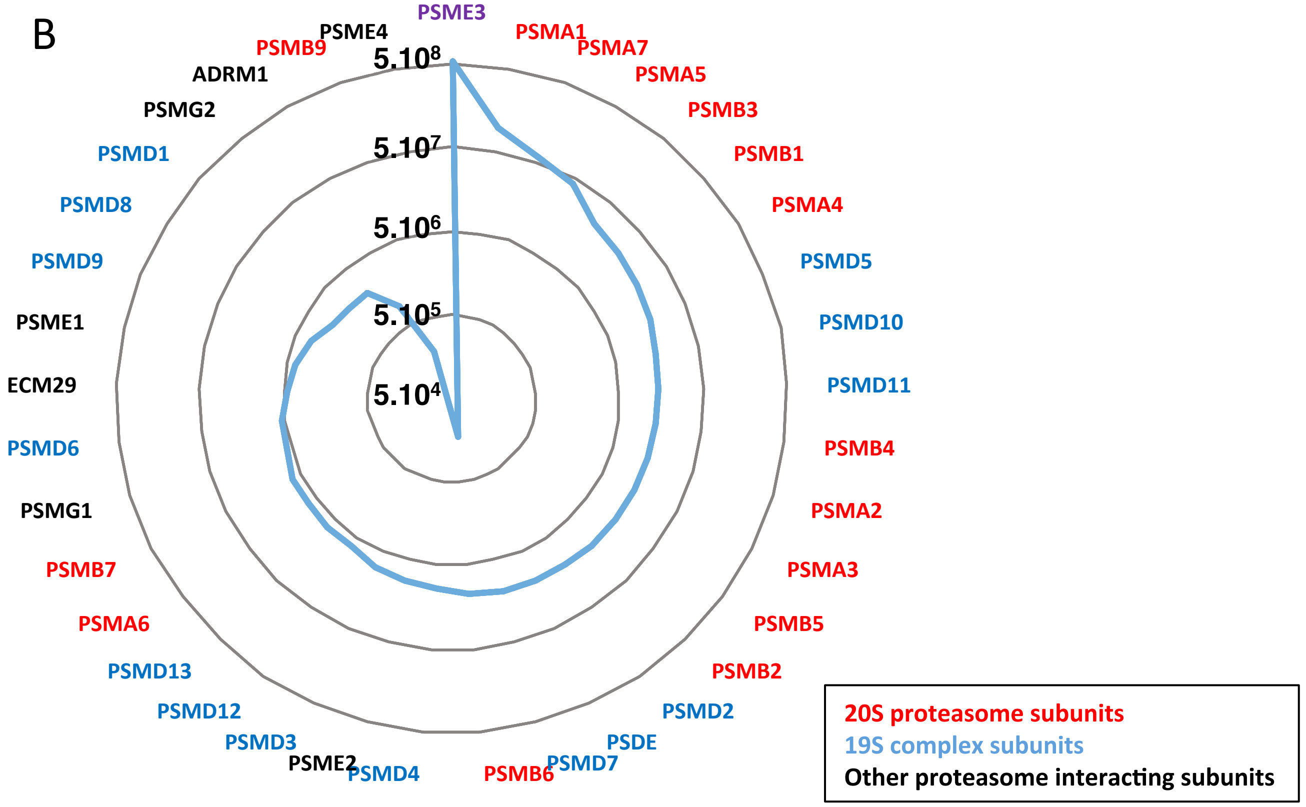

PSME3
PSMA1
PSMA7
PSMA5
PSMB3
PSMB1
PSMA4
PSMD5
PSMD10
PSMD11
PSMB4
PSMA2
PSMA3
PSMB5
PSMB2
PSMD2
PSDE
PSMD7
PSMB6
PSMD4
PSME2
PSMD3
PSMD12
PSMD13
PSMA6
PSMB7
PSMG1
PSMD6
ECM29
PSME1
PSMD9
PSMD8
PSMD1
PSMG2
ADRM1
PSMB9
PSME4
5.10^8
5.10^7
5.10^6
5.10^5
5.10^4
20S proteasome subunits
19S complex subunits
Other proteasome interacting subunits

*Figure S3*

A

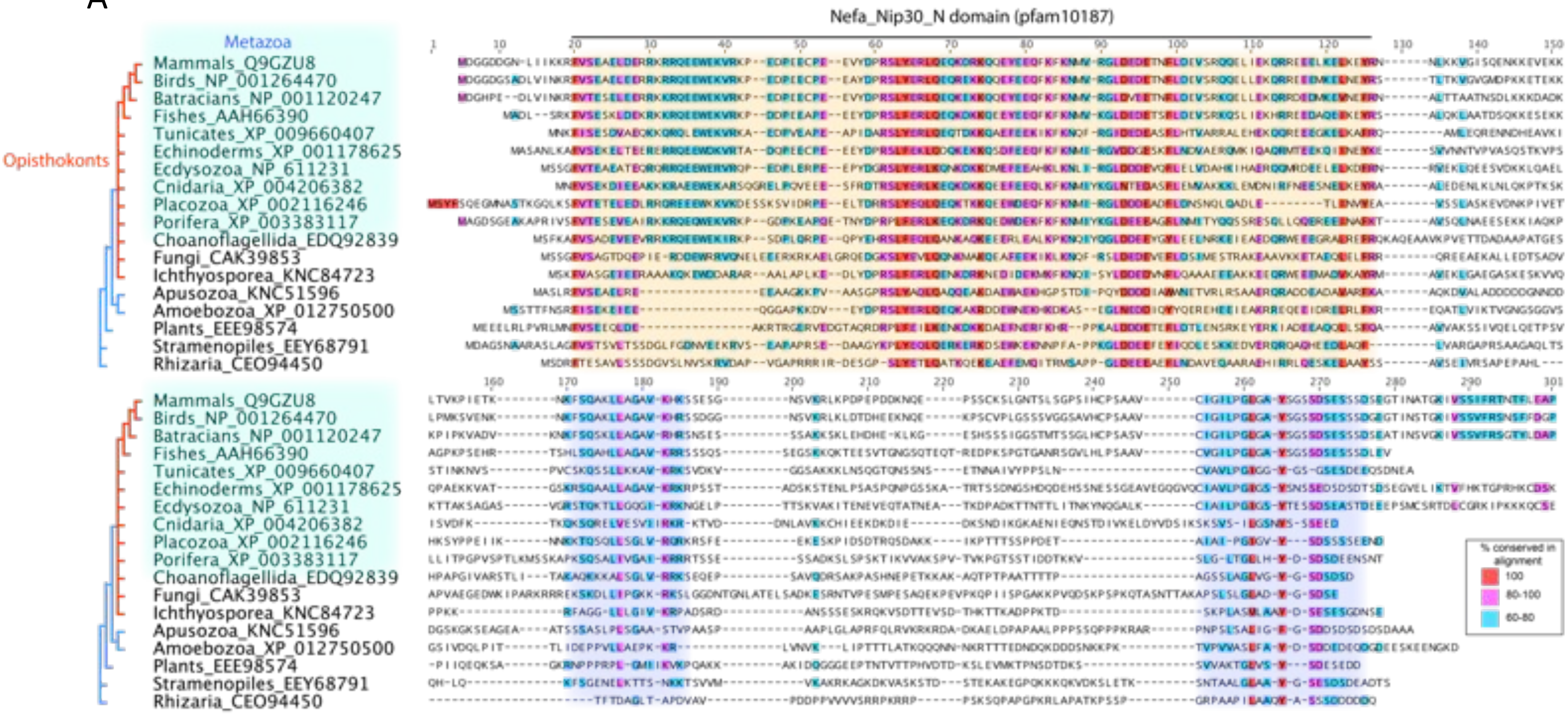
Nefa_Nip30_N domain (pfam10187)
Metazoa
Opisthokonts
Mammals_Q9GZU8
Birds_NP_001264470
Batracians_NP_001120247
Fishes_AAH66390
Tunicates_XP_009660407
Echinoderms_XP_001178625
Ecdysozoa_NP_611231
Cnidaria_XP_004206382
Placozoa_XP_002116246
Porifera_XP_003383117
Choanoflagellida_EDQ92839
Fungi_CAK39853
Ichthyosporea_KNC84723
Apusozoa_KNC51596
Amoebozoa_XP_012750500
Plants_EEE98574
Stramenopiles_EEY68791
Rhizaria_CEO94450
% conserved in alignment
100
80-100
60-80

*Figure S3*

B

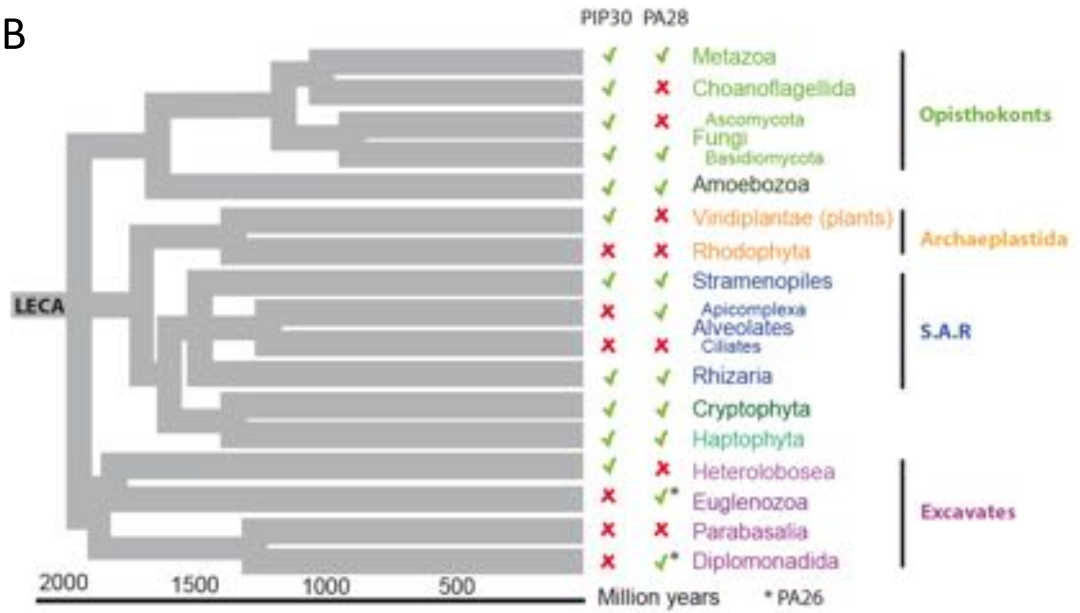
PIP30 PA28
LECA
Metazoa
Choanoflagellida
Ascomycota
Fungi
Basidiomycota
Amoebozoa
Viridiplantae (plants)
Rhodophyta
Stramenopiles
Apicomplexa
Alveolates
Ciliates
Rhizaria
Cryptophyta
Haptophyta
Heterolobosea
Euglenozoa
Parabasalia
Diplomonadida
Opisthokonts
Archaeplastida
S.A.R
Excavates
2000
1500
1000
500
Million years
* PA26

*Figure S4*

A

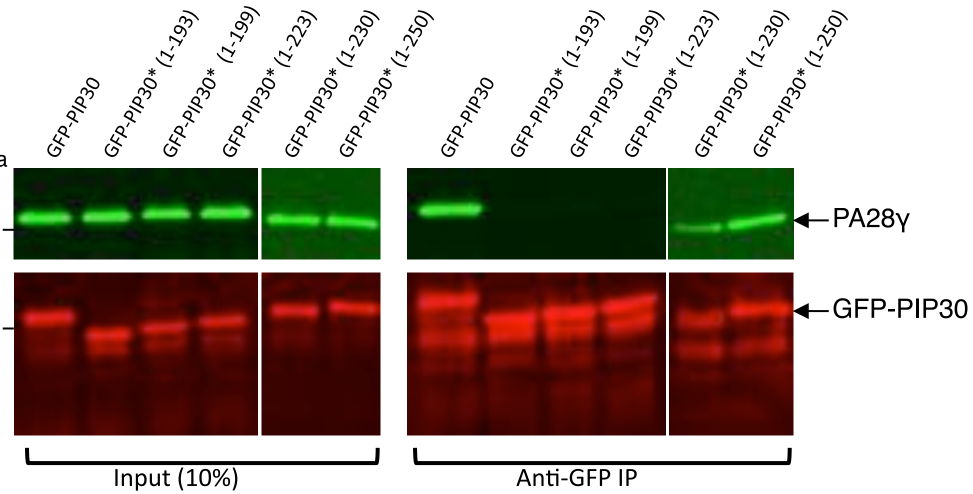


B

*CK2 consensus site*

**-S/T-** X- X- **E/D-**

| | |
|---|---|
| **WT** | 221- S**S$_{222}$DSE**SS**S$_{228}$DSE**- 231 |
| **SS-AA** | 221- S**A$_{222}$DSE**SS**A$_{228}$DSE**- 231 |
| **D$_{223}$E$_{225}$-KK** | 221- S**S$_{222}$KSK**SS**S$_{228}$DSE**- 231 |
| **D$_{229}$E$_{231}$-KK** | 221- S**S$_{222}$DSE**SS**S$_{228}$KSK**- 231 |

C

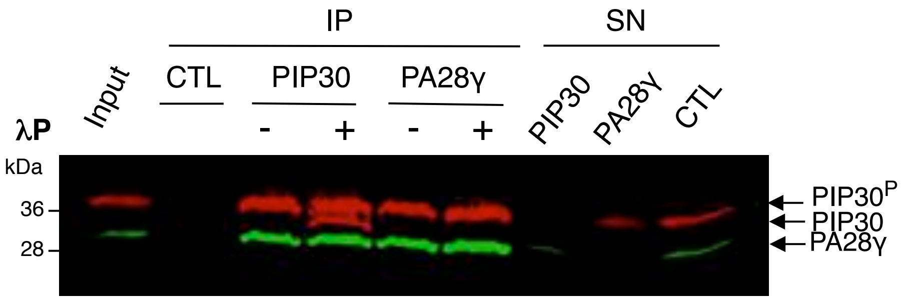

*Figure S5*

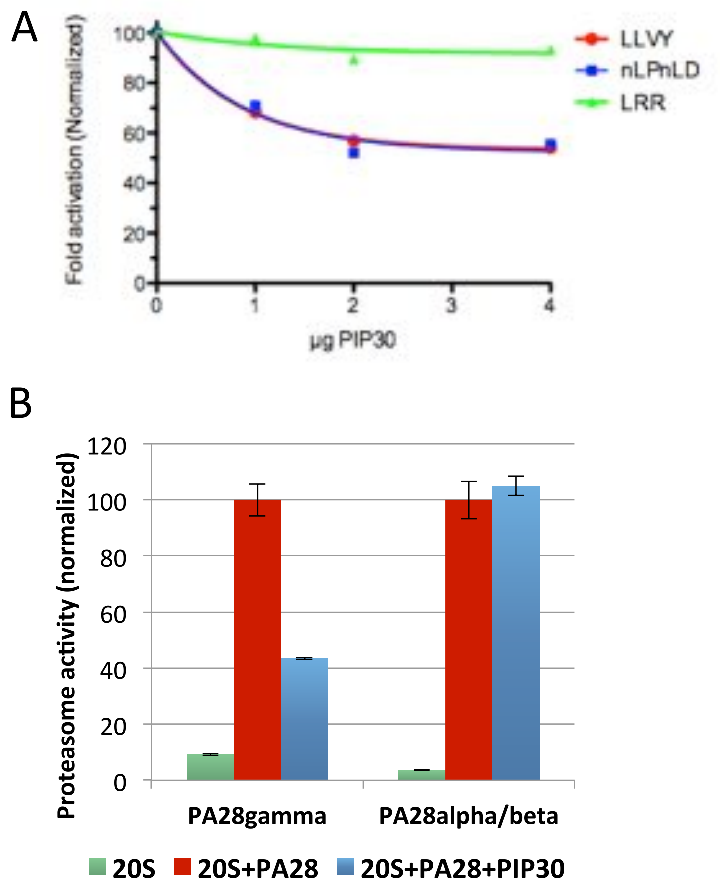
A
Fold activation (Normalized)
100
80
60
40
20
0
0
1
2
3
4
µg PIP30
LLVY
nLPnLD
LRR
B
Proteasome activity (normalized)
120
100
80
60
40
20
0
PA28gamma
PA28alpha/beta
20S
20S+PA28
20S+PA28+PIP30

*Figure S6*

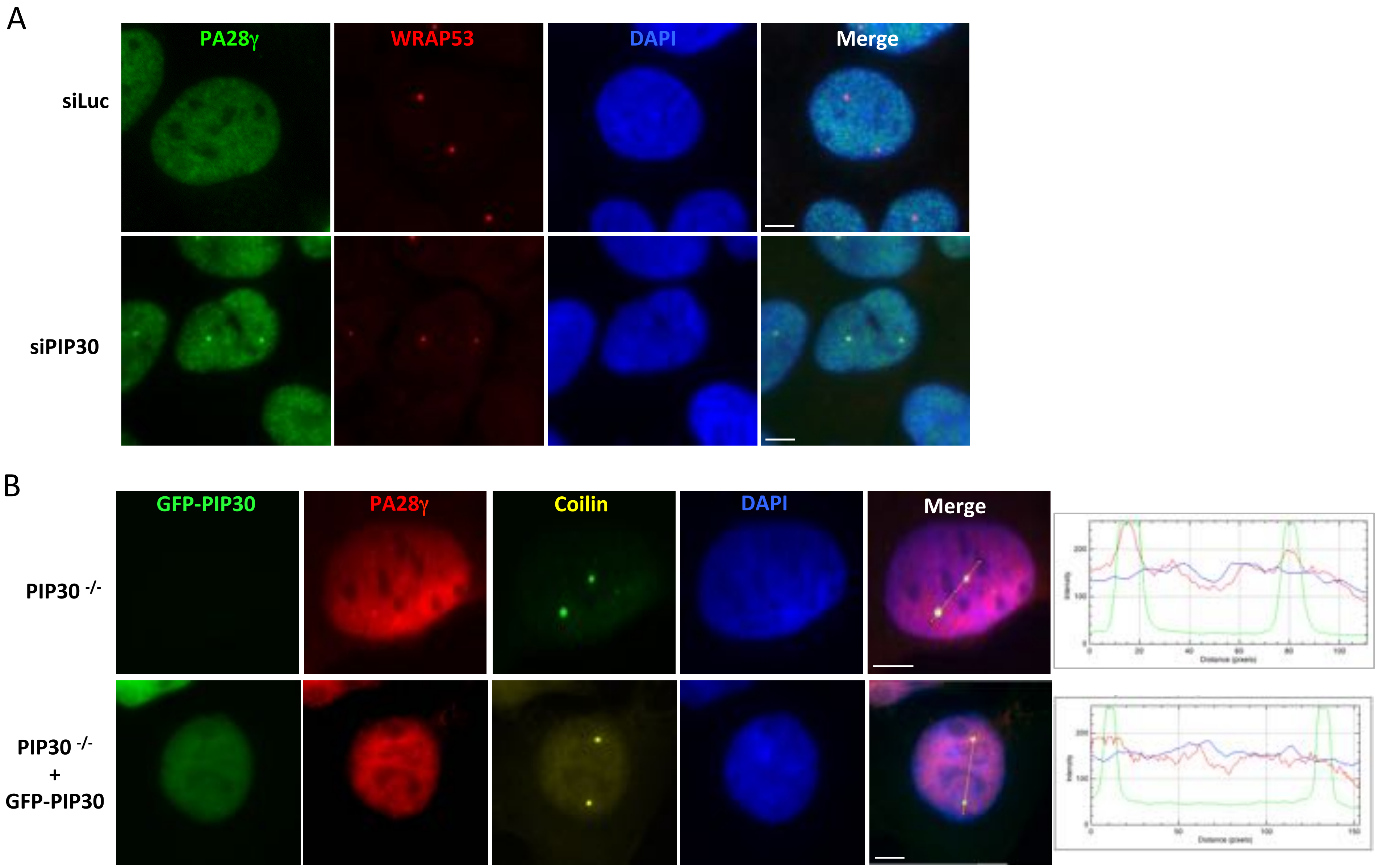
A
PA28γ
WRAP53
DAPI
Merge
siLuc
siPIP30
B
GFP-PIP30
PA28γ
Coilin
DAPI
Merge
PIP30 -/-
PIP30 -/-
+
GFP-PIP30

*Figure S7*

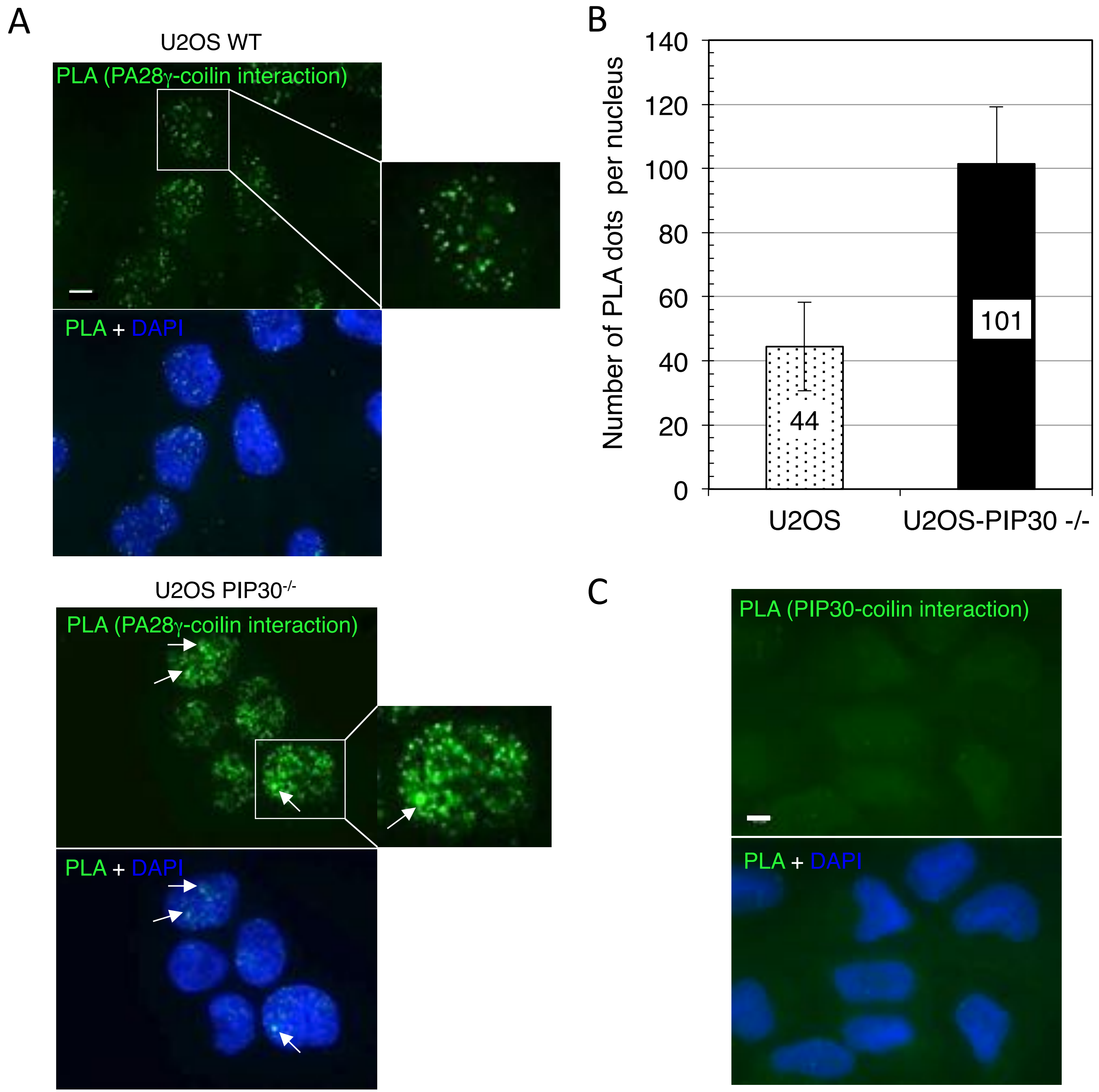

A
U2OS WT
PLA (PA28γ-coilin interaction)
PLA + DAPI
U2OS PIP30-/-
PLA (PA28γ-coilin interaction)
PLA + DAPI
B
Number of PLA dots per nucleus
140
120
100
80
60
40
20
0
44
101
U2OS
U2OS-PIP30 -/-
C
PLA (PIP30-coilin interaction)
PLA + DAPI

*Figure S8 (Characterization of tools)*

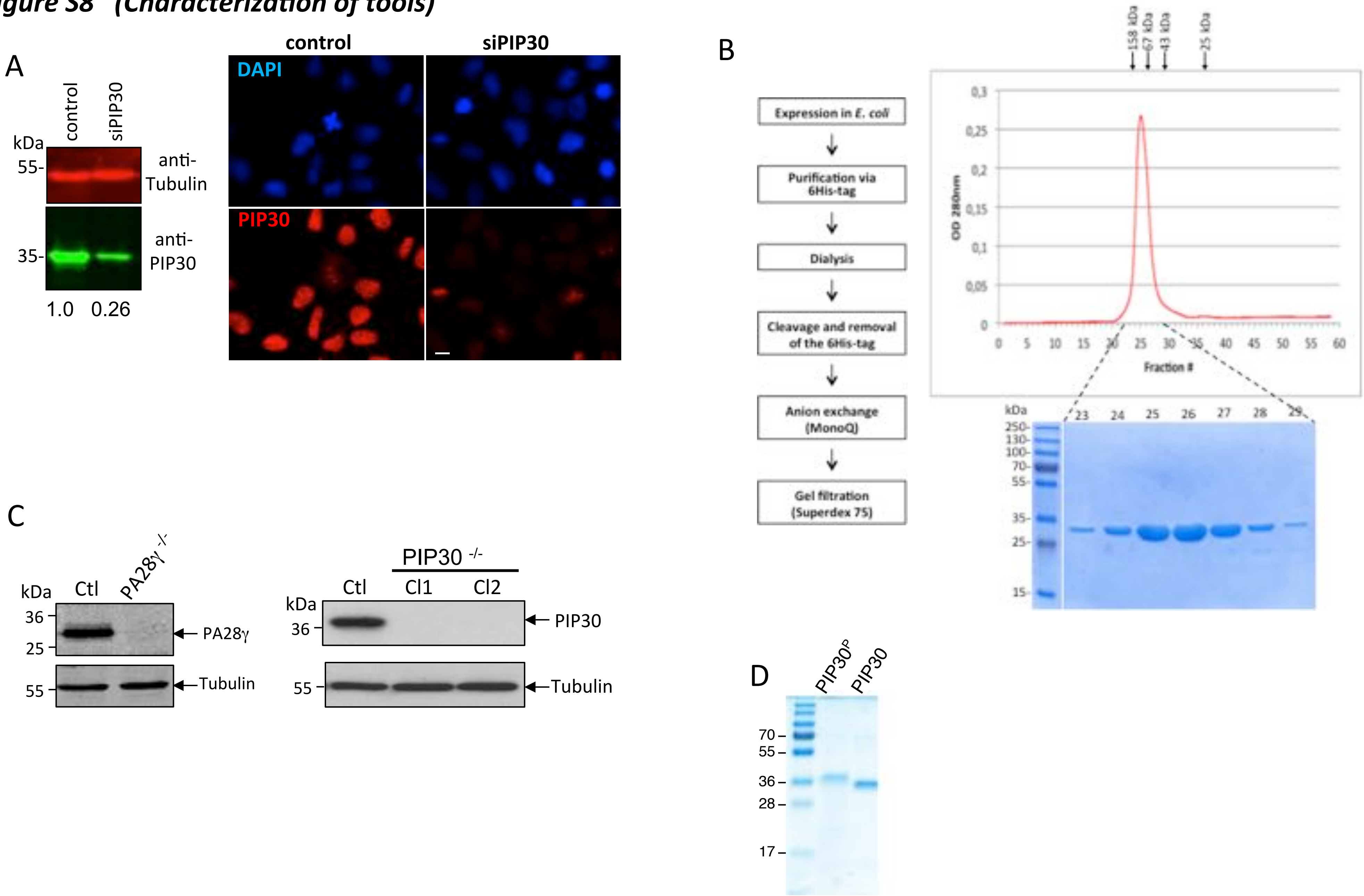

*Table S1*

| Gene Names | Protein IDs | Protein Names | Unique Peptides | Sequence Coverage [%] | Ratio H/L | Intensity H |
|---|---|---|---|---|---|---|
| PSMA1 | P25786 | Proteasome subunit alpha type-1 | 14 | 52,4 | 39,747 | 7888600 |
| PSME3 | P61289 | Proteasome activator complex subunit 3 | 25 | 76,8 | 14,914 | 2009700000 |
| CEP152 | O94986 | Centrosomal protein of 152 kDa | 3 | 2,5 | 12,05 | 3400400 |
| SMARCA4 | P51532 | SMARCA4 isoform 2 (SWI/SNF related, matrix associated, actin dependen | 16 | 20,1 | 6,2142 | 469900 |
| BRD9 | Q9H8M2 | Bromodomain-containing protein 9 | 8 | 17,8 | 4,394 | 338740 |
| NDUFS8 | O00217 | NADH dehydrogenase [ubiquinone] iron-sulfur protein 8, mitochondrial | 7 | 36,2 | 3,982 | 33060000 |
| MATR3 | P43243 | Matrin-3 | 2 | 5,1 | 3,7907 | 176150 |
| FAM192A | Q9GZU8 | NEFA-interacting nuclear protein NIP30 | 14 | 48 | 3,258 | 3433900 |
| FAU | P62861 | Ubiquitin-like protein FUBI;40S ribosomal protein S30 | 3 | 14,3 | 2,9776 | 86965 |
| HNRNPL;HNRPL | P14866 | Heterogeneous nuclear ribonucleoprotein L | 4 | 7,1 | 2,675 | 775540 |
| PLEC1 | Q15149 | Plectin-1;Hemidesmosomal protein 1;Plectin-11 | 3 | 68,4 | 2,5439 | 105370000 |
| U2AF2;U2AF65 | P26368 | Splicing factor U2AF 65 kDa subunit | 16 | 35,2 | 2,4359 | 5525600 |
| U2AF1;U2AF35 | Q01081 | Splicing factor U2AF 35 kDa subunit | 8 | 35 | 2,3661 | 5108200 |
| HNRNPM;HNRPM | P52272 | Heterogeneous nuclear ribonucleoprotein M | 6 | 9,7 | 2,3499 | 2570200 |
| SF3B3 | Q15393 | Splicing factor 3B subunit 3 | 8 | 8,3 | 2,3467 | 2011200 |
| HNRNPF;HNRPF | P52597 | Heterogeneous nuclear ribonucleoprotein F | 2 | 13,5 | 2,3449 | 5383200 |
| SF3B14 | Q9Y3B4 | Pre-mRNA branch site protein p14;SF3B 14 kDa subunit | 5 | 47,2 | 2,1821 | 183820 |
| BCLAF1 | Q9NYF8 | Bcl-2-associated transcription factor 1 | 6 | 6,1 | 2,1019 | 4912900 |
| RPS26 | P62854 | 40S ribosomal protein S26 | 4 | 37,4 | 2,0653 | 591760 |
| RPL22 | P35268 | 60S ribosomal protein L22 | 5 | 55,5 | 1,995 | 588410 |
| SF3B1;SAP155 | O75533 | Splicing factor 3B subunit 1 | 20 | 19,1 | 1,992 | 11332000 |
| RP9 | Q8TA86 | Retinitis pigmentosa 9 protein | 14 | 49,3 | 1,9651 | 16962000 |
| RBM39 | Q14498 | RNA-binding protein 39 | 22 | 52,1 | 1,9309 | 46370000 |
| ERH | P84090 | Enhancer of rudimentary homolog | 6 | 51 | 1,7589 | 5241600 |
| DHX9;DDX9 | Q08211 | ATP-dependent RNA helicase A;DEAH box protein 9 | 2 | 2,2 | 1,758 | 191250 |
| SFRS12IP1 | Q8N9Q2 | Protein SFRS12IP1;p18SRP | 4 | 21,4 | 1,726 | 1334100 |
| MMTAG2 | Q9BU76 | Multiple myeloma tumor-associated protein 2 | 6 | 22,8 | 1,7134 | 597760 |
| RPS27 | P42677 | 40S ribosomal protein S27 | 2 | 39,3 | 1,7133 | 368970 |
| ZCCHC17 | Q9NP64 | Nucleolar protein of 40 kDa | 4 | 21 | 1,7021 | 587080 |
| DHX15 | O43143 | DEAH box protein 15 | 4 | 5,8 | 1,6943 | 725370 |
| SNRPD2;SNRPD1 | P62316;A8K797 | Small nuclear ribonucleoprotein Sm D2 | 7 | 55,1 | 1,5949 | 312580 |
| DYNLL1 | P63167 | Dynein light chain 1, cytoplasmic | 2 | 20,2 | 1,589 | 320610 |
| DDX5 | P17844 | Probable ATP-dependent RNA helicase DDX5 | 13 | 30 | 1,5841 | 20552000 |
| THRAP3;TRAP150 | Q9Y2W1 | Thyroid hormone receptor-associated protein 3 | 13 | 14,9 | 1,5195 | 13683000 |
| PRPF40A | O75400 | Pre-mRNA-processing factor 40 homolog A | 5 | 25 | 1,4285 | 265810 |

*Table S2*

| Gene names | Protein IDs | Protein names | Unique peptides | Sequence coverage [%] | Ratio H/L | Intensity H |
|---|---|---|---|---|---|---|
| **GFP** | Q9U6Y5 | Green Fluorescent Protein | 11 | 53 | 44,917 | 15118000000 |
| UBB;RPS27A;UBC;UBA52;UBI | J3QS39;J3QTR3; | Ubiquitin-60S ribosomal protein L40;Ubiquitin;60S ribosomal protein L40 | 4 | 50,5 | 26,981 | 265340000 |
| **PSME3** | P61289 | Proteasome activator complex subunit 3 | 28 | 86,2 | 16,491 | 36430000000 |
| FAM192A;NIP30 | Q9GZU8 | Protein FAM192A | 12 | 63,5 | 15,779 | 182930000 |
| PSMA7;PSMA8 | O14818;Q8TAA | Proteasome subunit alpha type-7;Proteasome subunit alpha type-7-like | 2 | 12,9 | 3,7274 | 3118600 |
| COIL | P38432 | Coilin | 3 | 5,9 | 2,6545 | 3095400 |
| VDAC2 | P45880 | Voltage-dependent anion-selective channel protein 2 | 3 | 16,5 | 1,6517 | 6038600 |
| RCC2 | Q9P258 | Protein RCC2 | 2 | 5,2 | 1,5253 | 6408700 |
| CAPZA1 | P52907 | F-actin-capping protein subunit alpha-1 | 6 | 46,2 | 1,4971 | 53597000 |
| CAPZB | P47756 | F-actin-capping protein subunit beta | 4 | 17,3 | 1,4463 | 9878300 |
| KPNB1 | Q14974 | Importin subunit beta-1 | 2 | 3,2 | 1,4106 | 2802800 |

*Table S3*

| Gene names | Protein IDs | Protein names | Unique peptides | Sequence coverage [%] | Ratio H/L | Intensity H |
|---|---|---|---|---|---|---|
| GFP | Q9U6Y5 | Green Fluorescent Protein | 4 | 23,3 | 5,0732 | 101700000 |
| PSME3 | P61289 | Proteasome activator complex subunit 3 | 8 | 37,8 | 4,583 | 109590000 |
| FAM192A; NIP30 | Q9GZU8;Q6P4H7 | Protein FAM192A | 9 | 37 | 4,5533 | 69861000 |
| RPL22 | P35268 | 60S ribosomal protein L22 | 2 | 18,8 | 4,1196 | 533160000 |
| GEMIN5 | Q8TEQ6;Q58EZ8 | Gem-associated protein 5 | 2 | 1,4 | 2,8474 | 15548000 |

***Table S4***

| GN | AC | ID | Description | Best Id Score in Band # | Max Score | Coverage (%) | #QPep | T-Test | Enrichment Factor |
|---|---|---|---|---|---|---|---|---|---|
| ADRM1 | Q16186 | ADRM1_HUMAN | ADRM1_HUMAN Proteasomal ubiquitin receptor ADRM1 | 5 | 379 | 19,66 | 4 | 5,52E-04 | 23,5 |
| FAM192A | Q9GZU8 | F192A_HUMAN | F192A_HUMAN Protein FAM192A | 6 | 237 | 18,11 | 5 | 4,35E-04 | 6,39 |
| IDE | P14735 | IDE_HUMAN | IDE_HUMAN Insulin-degrading enzyme | 3 | 108 | 2,65 | 7 | 9,12E-04 | 6,99 |
| POMP | Q9Y244 | POMP_HUMAN | POMP_HUMAN Proteasome maturation protein | 10 | 39 | 7,09 | 1 | 4,54E-04 | 53,6 |
| PSMA1 | P25786 | PSA1_HUMAN | PSA1_HUMAN Proteasome subunit alpha type-1 | 7 | 1756 | 57,79 | 16 | 2,85E-04 | 53,4 |
| PSMA2 | P25787 | PSA2_HUMAN | PSA2_HUMAN Proteasome subunit alpha type-2 | 9 | 2652 | 44,02 | 16 | 1,21E-04 | 73,49 |
| PSMA3 | P25788 | PSA3_HUMAN | PSA3_HUMAN Proteasome subunit alpha type-3 | 8 | 2216 | 47,84 | 18 | 1,86E-04 | 46,85 |
| PSMA4 | P25789 | PSA4_HUMAN | PSA4_HUMAN Proteasome subunit alpha type-4 | 8 | 2637 | 59,77 | 15 | 8,75E-05 | 40,14 |
| PSMA5 | P28066 | PSA5_HUMAN | PSA5_HUMAN Proteasome subunit alpha type-5 | 8 | 1805 | 34,02 | 10 | 1,33E-04 | 42,45 |
| PSMA6 | P60900 | PSA6_HUMAN | PSA6_HUMAN Proteasome subunit alpha type-6 | 8 | 5958 | 34,96 | 14 | 1,00E-04 | 59,62 |
| PSMA7 | O14818 | PSA7_HUMAN | PSA7_HUMAN Proteasome subunit alpha type-7 | 8 | 3405 | 39,11 | 21 | 2,16E-04 | 58,07 |
| PSMB1 | P20618 | PSB1_HUMAN | PSB1_HUMAN Proteasome subunit beta type-1 | 9 | 3045 | 37,34 | 14 | 1,76E-04 | 80,73 |
| PSMB10 | P40306 | PSB10_HUMAN | PSB10_HUMAN Proteasome subunit beta type-10 | 8 | 882 | 30,77 | 7 | 2,43E-04 | 45,52 |
| PSMB11 | A5LHX3 | PSB11_HUMAN | PSB11_HUMAN Proteasome subunit beta type-11 | 9 | 102 | 4,67 | 1 | 1,08E-03 | 31,43 |
| PSMB2 | P49721 | PSB2_HUMAN | PSB2_HUMAN Proteasome subunit beta type-2 | 9 | 1537 | 37,81 | 15 | 9,07E-05 | 50,39 |
| PSMB3 | P49720 | PSB3_HUMAN | PSB3_HUMAN Proteasome subunit beta type-3 | 9 | 3233 | 62,93 | 8 | 7,15E-04 | 64,02 |
| PSMB4 | P28070 | PSB4_HUMAN | PSB4_HUMAN Proteasome subunit beta type-4 | 8 | 2834 | 53,03 | 5 | 5,32E-04 | 68,91 |
| PSMB5 | P28074 | PSB5_HUMAN | PSB5_HUMAN Proteasome subunit beta type-5 | 9 | 3147 | 57,41 | 14 | 3,11E-02 | 7,09 |
| PSMB6 | P28072 | PSB6_HUMAN | PSB6_HUMAN Proteasome subunit beta type-6 | 9 | 1826 | 17,57 | 6 | 1,97E-04 | 23,44 |
| PSMB7 | Q99436 | PSB7_HUMAN | PSB7_HUMAN Proteasome subunit beta type-7 | 8 | 889 | 38,27 | 5 | 1,67E-03 | 52,6 |
| PSMB8 | P28062 | PSB8_HUMAN | PSB8_HUMAN Proteasome subunit beta type-8 | 9 | 4105 | 28,99 | 12 | 4,01E-05 | 47,68 |
| PSMB9 | P28065 | PSB9_HUMAN | PSB9_HUMAN Proteasome subunit beta type-9 | 9 | 4948 | 26,94 | 8 | 2,57E-05 | 67,52 |
| PSMC1 | P62191 | PRS4_HUMAN | PRS4_HUMAN 26S protease regulatory subunit 4 | 4 | 2825 | 34,77 | 28 | 1,71E-03 | 54,09 |
| PSMC2 | P35998 | PRS7_HUMAN | PRS7_HUMAN 26S protease regulatory subunit 7 | 5 | 3188 | 32,56 | 33 | 5,20E-03 | 3,63 |
| PSMC3 | P17980 | PRS6A_HUMAN | PRS6A_HUMAN 26S protease regulatory subunit 6A | 5 | 4553 | 71,98 | 19 | 1,81E-05 | 37,76 |
| PSMC4 | P43686 | PRS6B_HUMAN | PRS6B_HUMAN 26S protease regulatory subunit 6B | 5 | 1928 | 23,68 | 37 | 9,25E-04 | 47,65 |
| PSMC5 | P62195 | PRS8_HUMAN | PRS8_HUMAN 26S protease regulatory subunit 8 | 5 | 6735 | 42,12 | 26 | 9,45E-04 | 56,9 |
| PSMC6 | P62333 | PRS10_HUMAN | PRS10_HUMAN 26S protease regulatory subunit 10B | 5 | 2692 | 77,12 | 28 | 8,92E-04 | 36,45 |
| PSMD1 | Q99460 | PSMD1_HUMAN | PSMD1_HUMAN 26S proteasome non-ATPase regulatory subunit 1 | 2 | 3142 | 38,51 | 35 | 2,60E-05 | 30,68 |
| PSMD11 | O00231 | PSD11_HUMAN | PSD11_HUMAN 26S proteasome non-ATPase regulatory subunit 11 | 5 | 2451 | 62,09 | 29 | 2,77E-05 | 32,74 |
| PSMD12 | O00232 | PSD12_HUMAN | PSD12_HUMAN 26S proteasome non-ATPase regulatory subunit 12 | 5 | 3159 | 29,39 | 25 | 4,50E-06 | 58,64 |
| PSMD13 | Q9UNM6 | PSD13_HUMAN | PSD13_HUMAN 26S proteasome non-ATPase regulatory subunit 13 | 6 | 1755 | 39,1 | 16 | 3,37E-05 | 35,64 |
| PSMD14 | O00487 | PSDE_HUMAN | PSDE_HUMAN 26S proteasome non-ATPase regulatory subunit 14 | 6 | 1850 | 46,45 | 4 | 1,44E-04 | 20,12 |
| PSMD2 | Q13200 | PSMD2_HUMAN | PSMD2_HUMAN 26S proteasome non-ATPase regulatory subunit 2 | 3 | 3894 | 51,1 | 38 | 2,76E-06 | 28,22 |
| PSMD3 | O43242 | PSMD3_HUMAN | PSMD3_HUMAN 26S proteasome non-ATPase regulatory subunit 3 | 4 | 3171 | 45,51 | 39 | 8,17E-04 | 29,04 |
| PSMD4 | P55036 | PSMD4_HUMAN | PSMD4_HUMAN 26S proteasome non-ATPase regulatory subunit 4 | 9 | 616 | 4,77 | 10 | 1,04E-06 | 16,08 |
| PSMD6 | Q15008 | PSMD6_HUMAN | PSMD6_HUMAN 26S proteasome non-ATPase regulatory subunit 6 | 5 | 1832 | 53,98 | 27 | 1,78E-04 | 80,25 |
| PSMD7 | P51665 | PSD7_HUMAN | PSD7_HUMAN 26S proteasome non-ATPase regulatory subunit 7 | 6 | 1867 | 47,22 | 14 | 7,23E-04 | 26,21 |
| PSMD8 | P48556 | PSMD8_HUMAN | PSMD8_HUMAN 26S proteasome non-ATPase regulatory subunit 8 | 8 | 578 | 8,57 | 10 | 8,68E-04 | 22,29 |
| PSME1 | Q06323 | PSME1_HUMAN | PSME1_HUMAN Proteasome activator complex subunit 1 | 8 | 1645 | 61,85 | 18 | 3,46E-04 | 40,42 |
| PSME2 | Q9UL46 | PSME2_HUMAN | PSME2_HUMAN Proteasome activator complex subunit 2 | 8 | 804 | 24,69 | 10 | 1,73E-04 | 15,71 |
| PSME3 | P61289 | PSME3_HUMAN | PSME3_HUMAN Proteasome activator complex subunit 3 | 7 | 654 | 44,88 | 17 | 4,94E-04 | 19,8 |
| PSME4 | Q14997 | PSME4_HUMAN | PSME4_HUMAN Proteasome activator complex subunit 4 | 2 | 188 | 3,36 | 5 | 3,84E-02 | 3,88 |
| PSMG1 | O95456 | PSMG1_HUMAN | PSMG1_HUMAN Proteasome assembly chaperone 1 | 7 | 154 | 21,53 | 8 | 1,85E-03 | 6,05 |
| RAD23A | P54725 | RD23A_HUMAN | RD23A_HUMAN UV excision repair protein RAD23 homolog A | 4 | 53 | 7,71 | 1 | 1,61E-02 | 2,89 |
| RAD23B | P54727 | RD23B_HUMAN | RD23B_HUMAN UV excision repair protein RAD23 homolog B | 4 | 249 | 36,43 | 10 | 1,33E-02 | 4,11 |
| SCCPDH | Q8NBX0 | SCPDH_HUMAN | SCPDH_HUMAN Probable saccharopine dehydrogenase | 5 | 105 | 6,06 | 2 | 4,92E-02 | 2,04 |
| TFG | Q92734 | TFG_HUMAN | TFG_HUMAN Protein TFG | 6 | 744 | 35,25 | 8 | 9,43E-03 | 2,91 |
| UBE3C | Q15386 | UBE3C_HUMAN | UBE3C_HUMAN Ubiquitin-protein ligase E3C | 2 | 474 | 13,39 | 14 | 5,70E-04 | 14,39 |
| UBLCP1 | Q8WVY7 | UBCP1_HUMAN | UBCP1_HUMAN Ubiquitin-like domain-containing CTD phosphatase 1 | 6 | 36 | 10,69 | 2 | 4,19E-02 | 2,29 |
| UBQLN2 | Q9UHD9 | UBQL2_HUMAN | UBQL2_HUMAN Ubiquilin-2 | 3 | 196 | 7,69 | 3 | 9,45E-03 | 4,27 |
| UCHL5 | Q9Y5K5 | UCHL5_HUMAN | UCHL5_HUMAN Ubiquitin carboxyl-terminal hydrolase isozyme L5 | 6 | 295 | 35,56 | 8 | 2,00E-03 | 2,73 |
| USP14 | P54578 | UBP14_HUMAN | UBP14_HUMAN Ubiquitin carboxyl-terminal hydrolase 14 | 4 | 1619 | 41,5 | 17 | 2,03E-03 | 5,87 |
